# Cell migration CRISPRi screens in human neutrophils reveal regulators of context-dependent migration and differentiation state

**DOI:** 10.1101/2022.12.16.520717

**Authors:** Nathan M. Belliveau, Matthew J. Footer, Emel Akdogan, Aaron P. van Loon, Sean R. Collins, Julie A. Theriot

## Abstract

Neutrophils are the most abundant leukocyte in humans and provide a critical early line of defense as part of our innate immune system. Their exquisite sensitivity to chemical gradients and ability to rapidly migrate make them especially suited to protect against infection. However, their terminal differentiation status and short lifetime (on the order of days) have hindered their study. Furthermore, while modern CRISPR-based gene perturbation strategies now allow comprehensive, genome-scale screens in human cells, their application to complex and dynamic processes like cell migration remain limited. Using HL-60 cells, a leukemia cell line that can be differentiated into neutrophil-like cells, we have developed multiple cell migration screen strategies that provide comprehensive, genome-wide discovery of molecular factors that are critical for directed (chemotaxis), undirected (chemokinesis), and 3D amoeboid cell migration in these fast-moving cells. Combining these assays with additional, pooled, genome-wide CRISPR interference dropout screens of cell proliferation and neutrophil differentiation, we have identified a comprehensive set of genes that are important across the processes of cellular growth, differentiation, and migration. This combined dataset highlights a particular reliance upon mTORC1 signaling that alters neutrophil lifetime, migration phenotype, and sensitivity to chemotactic cues. Across our cell migration screens, we identified several hundred genes important for migration including those with specific roles only in particular migratory contexts. This genome-wide screening strategy, therefore, provides an invaluable approach to the study of neutrophils and provides a resource that will inform future studies of cell migration in these and other rapidly migrating cells.

## INTRODUCTION

Among the cells of our immune system, neutrophils are the most abundant cell type and provide a vital early response in host defense by migrating to sites of infection or tissue wounding ^1, 2^. Paramount to their success is an exquisite sensitivity to chemical gradients, extremely rapid migration speeds on the order of 5 - 20 µm/min, and an ability to perform directed migration over long distances and through a wide variety of distinct tissue environments ^3–5^. To provide an effective response, neutrophils must constantly integrate physical and chemical cues as they move from the blood into tissue to reach the site of response ^6^. Although many cell types, including human neutrophils, can adapt to perform rapid migration in a wide variety of environments, relatively little is known about how specific molecular players may change or adapt as the cell migration context and environment change.

The emergence of CRISPR-based gene perturbation approaches and robust genome-wide targeted guide libraries now make it possible to perform unbiased functional genomics in human cells ^7–9^. These approaches offer significant technical improvements over past strategies such as RNAi and offer the opportunity to more comprehensively identify the genes involved in a biological process^10^. However, the use of genome-wide CRISPR-based screens to study complex and dynamic cellular processes has been more limited, with only a few notable exceptions where complex enrichment methods have been applied to identify factors important for phagocytosis and for cell motility ^11–14^. Extending these tools to the complex and varied behaviors of human cells through the development of new functional screening strategies is expected to provide new biological insights.

To extend these tools to perform a comprehensive screen of neutrophil cell migration, there are several notable challenges. First, their terminal differentiation status and short lifetime, on the order of days, limit the engineering and use of primary human cells^15^. Genome-wide screens also require millions of cells, demanding that the assays to assess the relevant biological response be relatively simple and easily scalable ^16^. For example, current pooled CRISPR strategies commonly use simple selections such as survival after a drug treatment, or enrichments of a cell population with fluorescence-activated cell sorting, to identify genetic perturbations of interest. In contrast, the study of cell migration has relied heavily on microscopy to assess behavior that is generally limited to tens or several hundred cells ^17, 18^. Furthermore, the context and physical environment is expected to alter the observed migratory behavior, highlighting the importance of assay design in the assessment and interpretation of such data.

In this work, we present the results of several pooled genome-wide CRISPRi screens that provide a comprehensive, genome-wide look at molecular factors contributing to distinct forms of cell migration, including directed migration (chemotaxis), undirected migration (chemokinesis), and 3D amoeboid migration through extracellular matrix. We use the acute promyelocytic leukemia HL-60 cell line ^19^, which can be differentiated in culture into rapidly migrating neutrophil-like cells. This choice of cell line allowed us to also perform conventional, pooled genome-wide CRISPRi dropout screens of proliferation and differentiation, providing several additional dimensions to assess and distinguish biological function for this important cell type. We confirm known molecular mechanisms contributing to cell proliferation and differentiation and identify unexpected mechanisms that alter neutrophil differentiation, survival, and cell migration. We find a near-perfect correlation between the genes important for chemotaxis and chemokinesis, suggesting that both modes of migration are mechanistically identical. Lastly, we use the results from our different screens of cell migration to distinguish between adhesion-dependent and independent cell migration, ultimately identifying several hundred genes that are important across these different migratory contexts. This work demonstrates an invaluable strategy to study cell migration and provides a resource that will apply to future studies of migration in neutrophils and other rapidly migrating cell types.

## RESULTS

### Pooled CRISPRi screens identify genes that alter cell proliferation, neutrophil differentiation, and cell migration

To identify novel regulators important for neutrophil biology, as well as facilitate our primary goal to identify genetic factors critical for cell migration, we used the immortalized HL-60 cell line human tissue culture line derived from a patient with acute promyelocytic leukemia, which resemble progenitor cells along the normal granulocyte-monocyte lineage ^2, 15, 20^. The proliferating, undifferentiated HL-60 cells (uHL-60) can be induced to differentiate into a neutrophil-like cell type (dHL-60) by treatment with the signaling molecule all-trans retinoic acid (ATRA) or with the organic solvent dimethylsufoxide (DMSO) ^19^. After differentiation, the chemotactic and migratory behavior of dHL-60 cells closely mimic those of primary neutrophils, and they are able to clear fungal infections in neutropenic mice ^21–23^.

To achieve reliable gene knockdown in uHL-60 cells, we used dCas9-KRAB linked by a proteolysis-resistant 80 amino acid XTEN linker ^24, 25^, driven by an EF1ɑ promoter that was placed downstream of a minimal-ubiquitous chromatin opening (UCOE) element to prevent gene silencing ^24, 26^. We validated the efficiency of knockdown in this cell line by targeting the CD4 gene, with immunofluorescence flow cytometry measurements demonstrating robust knockdown using this construct (Fig. 1A), which was substantially better than the efficacy of several other dCas9-KRAB constructs with the same target (Fig. S1).

**Figure 1:**
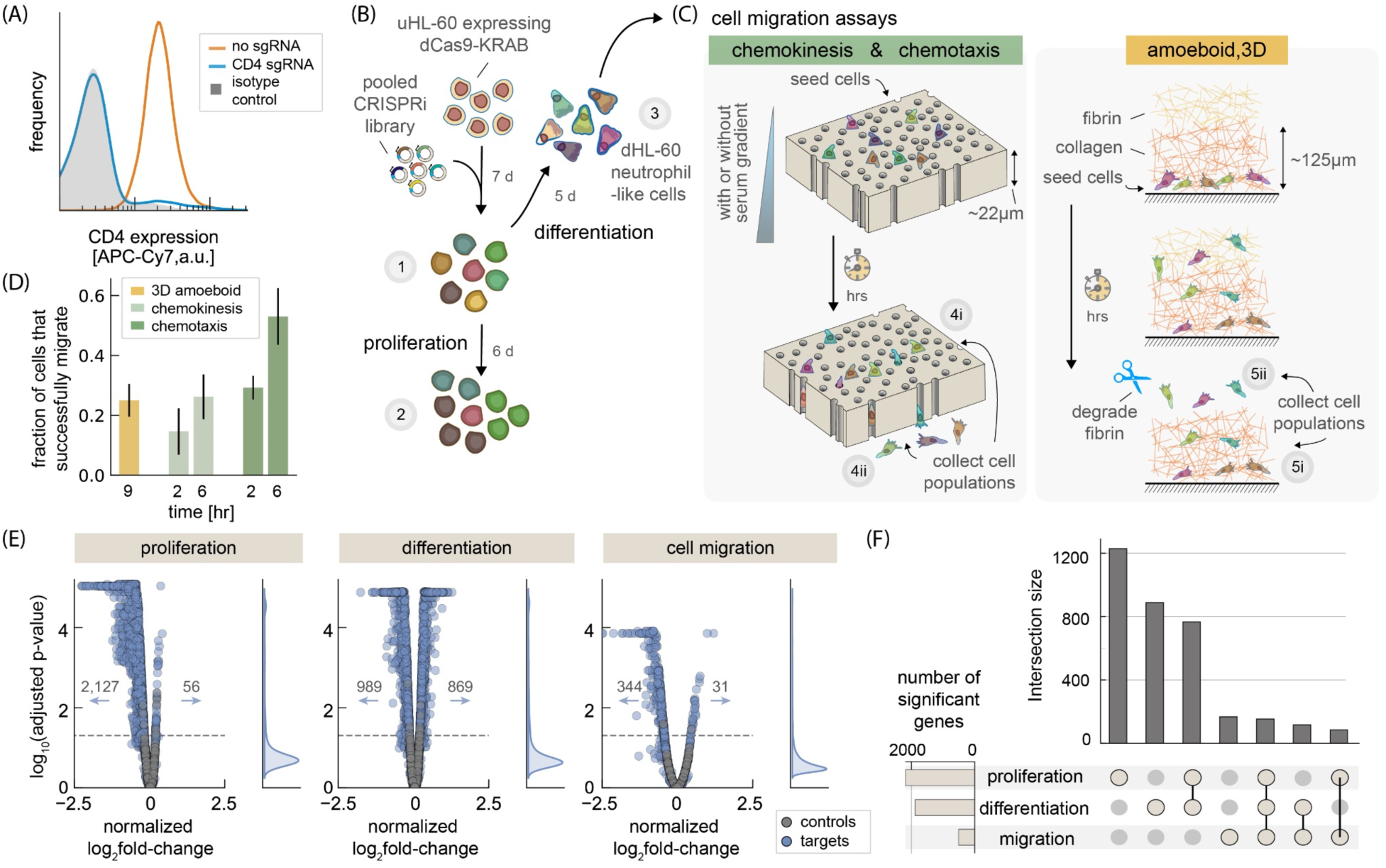
Genome-wide CRISPRi screens of proliferation, differentiation, and cell migration. (A) CRISPRi gene knockdown using sgRNA targeting the CD4 gene in uHL-60 cells. Flow cytometry immunofluorescence of CD4 protein expression shows near-complete loss of protein (blue) when compared to isotype antibody control (gray, shaded), representing cellular autofluorescence. Normal expression of CD4 in uHL-60 cells is shown in orange. (B) Schematic of pooled CRISPRi dropout experiments of proliferation of uHL-60 cells and differentiation into dHL-60 neutrophils. A pooled sgRNA library was integrated into uHL-60 cells expressing dCas9-KRAB using lentiviral transduction and selected for stable integration with puromycin for seven days. Proliferation was assayed by comparing sgRNA abundances following six days of growth (∼24 hr doubling time) (set 2 versus set 1). Differentiation was assayed by comparing sgRNA distribution in dHL-60 neutrophils with uHL-60 cells (set 3 versus set 1). The proliferation screen was performed four times, while the differentiation screen was performed eight times. (C) Schematic of pooled CRISPRi cell migration assays. Migration in dHL-60 cells was assayed across three experiments: The first two assessed migration through track-etch membranes with 3 *μm* diameter pores, stimulating migration either through the presence of a serum gradient (chemotaxis, 10% hiFBS added to the bottom reservoir) or in a uniform serum environment (chemokinesis, uniform 10% hiFBS added throughout media). The third assay assessed migration through an extracellular matrix composed of collagen and fibrin (see *Methods* for additional details). For quantification of migration screens, sgRNA abundance in both migratory fractions (sets 4i and 5i) and remaining cells (sets 4ii and 5ii) were compared to our initial dHL-60 library (set 3). Membrane, pores and cells drawn to scale. (D) Quantification of the migratory cell populations collected in the different migratory screens (i.e. sets 4ii and 5ii). Migration through 3 *μm* diameter pores of track-etch membranes was assayed at two time points (2 hr, 6 hr) and (+/-) a gradient of hiFBS. Cells were collected from the extracellular matrix after nine hours. Error bars represent standard deviation (amoeboid, 3D: N = 6; chemokinesis: 2 hr, N = 16, 6 hr, N = 12; chemotaxis: 2 hr and 6 hr, N = 4). (E) Volcano plots showing the statistical significance across the CRISPRi screens of proliferation, differentiation, and cell migration. Data points represent the average log_2_ fold-change across the three sgRNA per gene. Cell migration values represent an average across all migration assays. Control data points were generated by randomly selecting groups of three control sgRNAs. Adjusted p-values were calculated using permutation tests with the dashed line representing a value of 0.05. See *Methods* for additional details on data analysis and number of replicates for each experiment. (F) Histogram plots show the number of significant genes using an adjusted p-value cutoff of 0.05 (left horizontal bar plot) and the intersection of genes across each of the screens (vertical bar plot). The dot diagram identifies the specific screens considered for each intersection in the vertical bar plot.

To identify genes regulating proliferation and differentiation, we performed dropout-type assays using a pooled genome-wide CRISPRi library (3 sgRNA per gene; Sanson et al., 2018) (Fig. 1B). Here, we identified genes whose knockdown perturb these processes by quantifying changes in sgRNA abundance within the pooled library, as compared to the appropriate starting population. In the case of proliferation, we quantified log_2_ fold-changes in sgRNA abundance in the uHL-60 cells following six days of growth, as compared to the abundance at day zero (immediately following selection for stable integration of the sgRNA). At a false discovery rate (fdr) of 0.05, we identified 2,127 genes that disrupted growth and only 56 genes whose knockdown appeared to enhance growth (see *Methods* for additional details). The genes we identified were generally well-correlated with those reported by Sanson *et al.*, who originally performed a drop-out proliferation screen in HT29 (colorectal adenocarcinoma) and A375 (melanoma) cell lines (Fig. S2, Pearson correlation coefficient, ρ = 0.6). These results gave us confidence that our genome-wide CRISPRi knockdown approach in uHL-60 cells was working as expected.

We induced differentiation to derive dHL-60 cells by incubating uHL-60 cells with 1.57% DMSO for five days ^19, 27, 28^. Comparing sgRNA abundance between the dHL-60 cells and their uHL-60 predecessors, we identified 989 genes that were depleted and 869 genes that were enriched (Fig. 1E). This ratio was strikingly different from our observation with proliferation, where only ∼2% of knockdowns led to an enrichment of sgRNAs. Since wild-type dHL-60 cells undergo terminal differentiation without further cell division ^15, 19^, we hypothesize that the enriched sgRNAs identified in our differentiation screen may follow from alterations to the initiation of differentiation and/or a change in the lifetime of differentiated cells, and look at these genes more closely in the next section.

To assay cell migration, we performed three independent migration experiments to assay contextually different aspects of cell migration. The first two involved migration through 3 µm diameter pores using track-etch membranes (Fig. 1C, left panel). Such membranes are commonly employed in chemotaxis transwell assays and provide a simplified mimic of migration by neutrophils through tight cellular junctions during transmigration across an endothelial layer ^17, 29^. Cells are placed above the membrane and a diffusible chemoattractant is added to a second reservoir below. Following diffusion, a gradient in chemoattractant concentration entices cells to migrate down, through the membrane. Here we used heat-inactivated fetal bovine serum (hiFBS) as a general stimulant of cell migration, and included 10% hiFBS in the bottom reservoir to probe chemotaxis. Absent any directional cue, neutrophils can also be stimulated to migrate in random directions following uniform chemical stimulation, in a process known as chemokinesis ^30, 31^. In order to determine whether any gene knockdowns would have specific effects on gradient-directed versus random cell migration in the context of chemical stimulation, we also performed chemokinesis assays by stimulating migration with 10% hiFBS in both reservoirs. For both the chemotaxis and chemokinesis assays, cells were collected following two-hour and six-hour time points (Fig. 1D). To assess migratory success, we separately collected both the migratory cells that made it to the bottom reservoir and the cells that remained above the track-etch membrane, calculating normalized log_2_ fold-change by comparing sgRNA abundances in these cell pools relative to a reference pool of dHL-60 cells (see *Methods* for further details).

Our final migration assay focused on probing the amoeboid three-dimensional (3D) migration characteristics of neutrophils by embedding cells in a synthetic extracellular matrix (ECM), in order to mimic migration through the intercellular spaces in tissue. Here we began by embedding cells at the bottom of a thin layer (∼200 µm) of collagen ECM that cells would need to traverse, prior to reaching a second layer of fibrin ECM where they would be recovered (Fig. 1C, right panel). To a first approximation, we expect cells to perform a random walk, whose mean-square displacement should scale with √*t* where *t* is the time cells migrate ^32^, requiring substantially more time for migration than through the thin track-etch membranes. We therefore only considered a longer, nine-hour time point prior to cell collection in these experiments. To collect the most migratory dHL-60 cells, we used the enzyme nattokinase, which has protease activity specific to fibrin ^33^, to degrade the upper fibrin layer. Separately, we collected the cells still embedded in collagen, again calculating normalized log_2_ fold-change by comparing sgRNA abundances in these cell sub-populations, relative to a reference population of dHL-60 cells.

Across our entire set of cell migration assays, we identified 344 genes that reduced the fraction of migratory cells and 31 genes that increased this fraction, relative to migration of control sgRNAs. While we look into the individual migration screens further in later sections, we were also interested in the overlap of identified genes across our assays of proliferation, differentiation, and cell migration. Inevitably, we would expect genes that perturb neutrophil differentiation to also diminish the ability of cells to migrate or perform other cellular functions specific to neutrophils. We found that nearly half of the genes identified in each screen of proliferation and differentiation were unique to those screens (Fig. 1F). Interestingly, while there was substantial overlap across all screens, only 116 genes were unique to differentiation and cell migration, with another 167 genes unique to cell migration.

### CRISPRi screens identify genes important for differentiation into migratory neutrophils

Stimulating the differentiation of leukemia cells using pharmacological agents remains a key strategy for the clinical treatment of acute promyelocytic leukemia ^34^. We were therefore interested in identifying what gene perturbations influenced the differentiation of uHL-60 cells into migratory, terminally differentiated (non-proliferating) dHL-60 cells. Here, we began by performing gene set enrichment analysis (GSEA) ^35^ to identify the pathways and gene categories that were overrepresented among the differentiation screen data (Fig. 2A). Among the most positively enriched gene sets, there was a positive enrichment across various metabolic processes, particularly genes involved in oxidative phosphorylation (electron transport chain, mitochondrial protein synthesis). We also found a depletion of genes associated with granulopoiesis, suggesting that our screen data is identifying genes specific to neutrophil differentiation.

**Figure 2:**
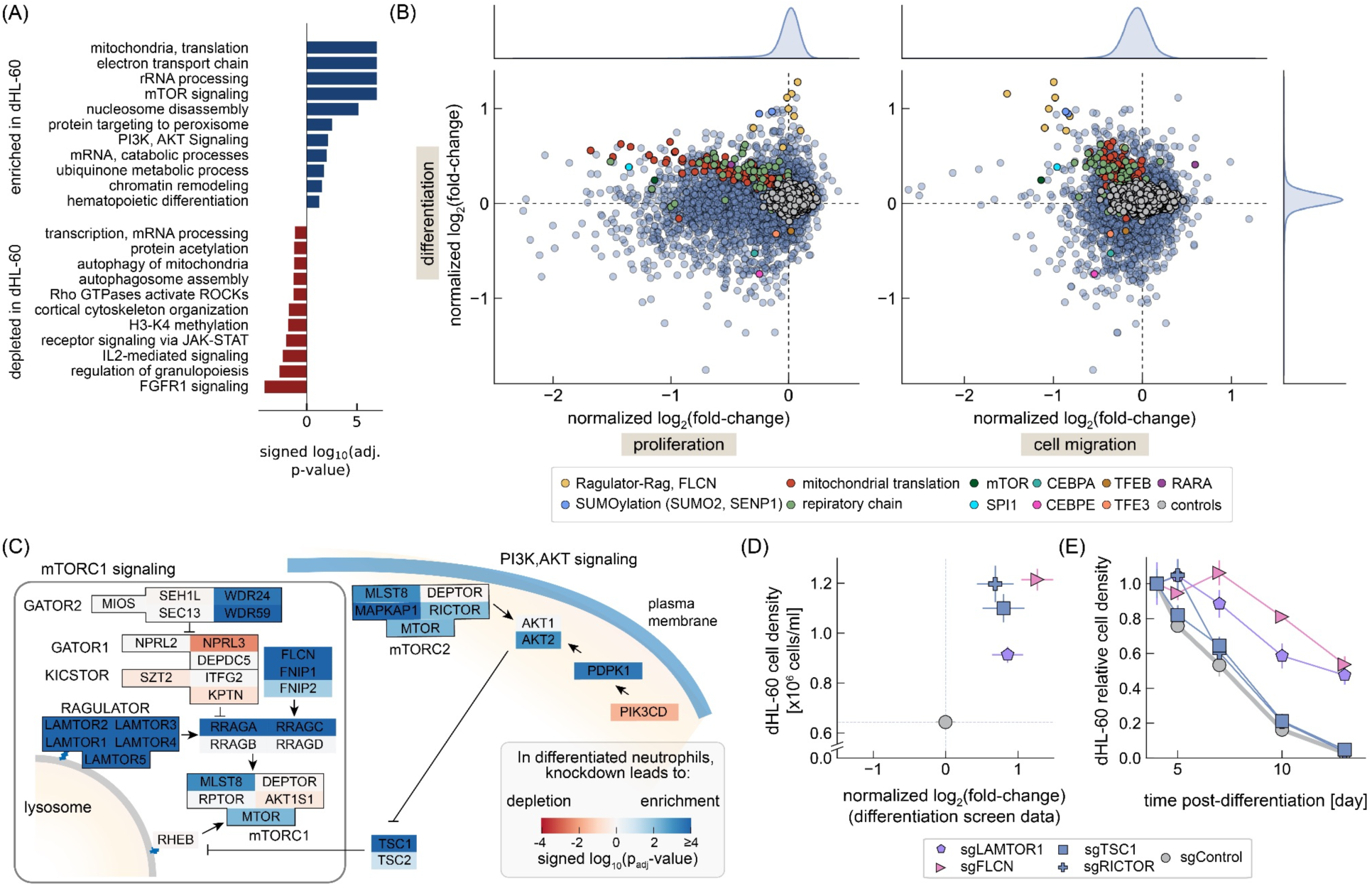
Identification of genes and pathways important for neutrophil differentiation. (A) Pathways enriched in our CRISPRi differentiation screen (dHL-60 cells relative to uHL-60 cells). Pathways that were associated with genes whose knockdown predominantly led to an enrichment of target sgRNAs in the dHL-60 cells are identified in blue, while those that decreased in abundance are in red. (B) Comparison of log_2_ fold-changes across the CRISPRi screens of proliferation, differentiation, and cell migration. Several gene sets identified through our pathway or known regulators of neutrophil differentiation are identified. (C) Schematic of mTORC1/ mTORC2 signaling pathway, color coded by signed statistical significance values (log_10_ p_adj_-value) from the differentiation screen results. Blue indicates gene targets whose sgRNA were enriched in the dHL-60 cells, while red indicates those that were depleted. (D) Differentiation screen results were confirmed across a number of sgRNA targets. Cell density was monitored at 5-days following the initiation of neutrophil differentiation. The dashed lines represent the values obtained for dHL-60 cells with a control sgRNA. Error bars represent standard deviation across replicates (N=8 for screen, N=3 cell density). (E) Change in dHL-60 cell density following differentiation. Note that media was replenished every three days, with cell density measurements corrected to account for changes in media volume and evaporation. Cell density measurements were normalized to day 4, following initiation of cell differentiation of uHL-60 cells. Error bars represent standard deviation across replicates (N=3).

To further distinguish genes whose knockdown specifically affected differentiation *per se* from those that generally perturbed basic cellular processes, we also plotted the differentiation log_2_ fold-change values against both our screens of proliferation and migratory data (Fig. 2B). Strikingly, knockdown of genes associated with oxidative phosphorylation and mitochondrial translation showed systematic effects in all three screens, with sgRNAs enriched following differentiation while inhibitory to both proliferation and migration (Figure 2B, red and green points). In contrast, sgRNAs targeting genes associated with mammalian target of rapamycin (mTOR) signaling were enriched following cell differentiation and depleted in the migration assays, but knockdown of these genes had no consequence on proliferation (Fig. 2B, yellow points). Interestingly, several genes involved in SUMOylation (SUMO2, SENP1), referring to the covalent attachment of small ubiquitin-like modifiers that alter protein function, also exhibited a similar pattern across the three screens as the mTOR-related genes. Protein SUMOylation has been shown to alter mTOR function directly and indirectly ^36, 37^, consistent with the possibility that these perturbations may also relate to mTOR signaling in the context of neutrophil differentiation, though we do not explore this further in this work.

We were also interested in whether regulatory genes known to be important for myeloid cell differentiation were represented in our data. PU.1 (SPI1), a transcription factor which is highly expressed in neutrophils and associated with terminal differentiation ^38, 39^, was identified across all screens and was among the strongest perturbations to migration (normalized log_2_ fold-change of -1.2, Fig. 2B right panel). Knockdown of the CCAAT/enhancer binding proteins C/EBPɑ (CEBPA) and C/EBP*ε* (CEBPE) resulted in a depletion of these cells in the dHL-60 CRISPRi library cells and also negatively perturbed migration. This depletion following knockdown of C/EBPε is consistent with recent experiments in mice, where its loss attenuated the progression of cells into mature neutrophils ^2^. Although we used DMSO (rather than all-*trans* retinoic acid, or ATRA) as the trigger for differentiation in our experimental protocol, nevertheless we found the retinoic acid receptor α (RARA) as one of the genes whose knockdown results in enrichment in the cell pool after differentiation, and these cells also were among the few where we observed enhanced migration in our chemotaxis and chemokinesis screens (normalized log_2_ fold-change of 0.7, Supplemental Data Table 3/4; Fig. 2B right panel). RARɑ is a DNA-binding protein that represses transcription of target genes in the absence of ligand ^40^, so it is likely that knockdown of the RARA gene is in some ways functionally equivalent to addition of ATRA. Notably, patients with acute promyelocytic leukemia, characterized by accumulation of undifferentiated neutrophil precursors, may be treated with ATRA as a way of inducing terminal differentiation in the cancerous cells ^41^.

Considering our differentiation screen results more broadly, we found that many of the identified genes that altered the course of HL-60 cell differentiation were also enriched in recent genome-scale efforts to characterize mouse embryonic stem cell differentiation ^42, 43^. Strikingly, in the roughly 500 genes reported as important for exit from pluripotency in mouse embryonic stem cells, half were present in our differentiation data set (adjusted p-value < 0.01 when applying gene set enrichment analysis, Fig. S2B). While those efforts represent a distinctly different developmental cell stage, this commonality suggests shared processes that may be important as mammalian cells change their proliferative status and undergo state transitions in differentiation.

### Disruption of folliculin and Ragulator-Rag signaling pathways potentiate differentiation phenotype and survival of dHL-60 neutrophils

Given the enrichment of sgRNAs associated with mTOR signaling during differentiation and migration, we wanted to further understand the consequence of these gene perturbations on cellular function. In humans, the protein kinase mTOR is a component of two distinct complexes, mTORC1, and mTORC2. While mTORC2 has previously been implicated in cell migration and chemotaxis ^44–48^, we were surprised to find significant effects of many genes associated with mTORC1 signaling as well (Fig. 2C). mTORC1 generally coordinates growth through its activity on the surface of lysosomes by mediating cellular changes in translation regulation, metabolism, and autophagy. The genes most enriched in the differentiation screen were directly upstream of mTORC1 and included the Rag guanosine triphosphatase (GTPase) A/B:C/D heterodimer, which recruits mTORC1 to the lysosomal membrane ^49^ via binding to the Ragulator complex (LAMTOR1-5) (Fig. 2C). Knockdown of folliculin (FLCN), a GTPase-activating protein that targets RagC/D and promotes an active state of the Rag heterodimer ^50^, led to a similar enrichment among dHL-60 cells in the differentiation screen and depletion in the migration screen.

We were interested in understanding how disruption of these signaling pathways altered differentiation, and began by characterizing in more detail how they might affect the progression of cells from uHL-60 into dHL-60. To this end, we generated individual stable cell lines expressing mTORC1-related sgRNAs targeting LAMTOR1, FLCN, TSC1, and also a cell line expressing an sgRNA targeting RICTOR, a key subunit of the mTORC2 complex. Following the initiation of neutrophil differentiation, all cell lines showed a higher cell density compared to a control sgRNA cell line, consistent with their overrepresentation in our differentiation screen results (Fig. 2D). For normal uHL-60 cells induced to differentiate by DMSO, cell densities stopped increasing by about 4 days after initial DMSO exposure and began to decline, presumably due to apoptosis of the terminally differentiated dHL-60s (Fig. 2E). Interestingly, LAMTOR1 and FLCN knockdown lines showed a distinct increase in cell lifetime by this assay (Fig. 2E, pink and purple symbols). Visually, these cells appeared healthier than our control sgRNA cell line and exhibited polarized neutrophil-like morphologies well past the standard 5-day time frame for generating dHL-60 cells used in our migration assays (see below).

We next assessed key molecular markers of neutrophil differentiation. Here we used flow cytometry to measure the induced surface expression of CD11b, an early differentiation marker also known as integrin *α*_M_ (ITGAM), and the fMLP receptor (FPR1) that recognizes chemoattractant N-formylated peptides ^51^. Both markers showed little to no expression in uHL-60 cells but were strongly induced in our dHL-60 cells (Fig. 3A). As positive controls for disruption of neutrophil differentiation, we constructed stable cell lines expressing sgRNAs targeting SPI1 and CEBPE. To assess induction of differentiation markers, we used principal component analysis to quantify the axis associated with co-induction of the two surface markers ITGAM and FPR1 (principal mode 1, Fig. 3B). As expected, sgRNAs targeting SPI1 and CEBPE reduced their expression. In contrast, we found that stable cell lines expressing sgRNAs targeting LAMTOR1 and FLCN exhibited higher induction of the differentiation markers (Fig. 3C, Fig. S3A).

**Figure 3:**
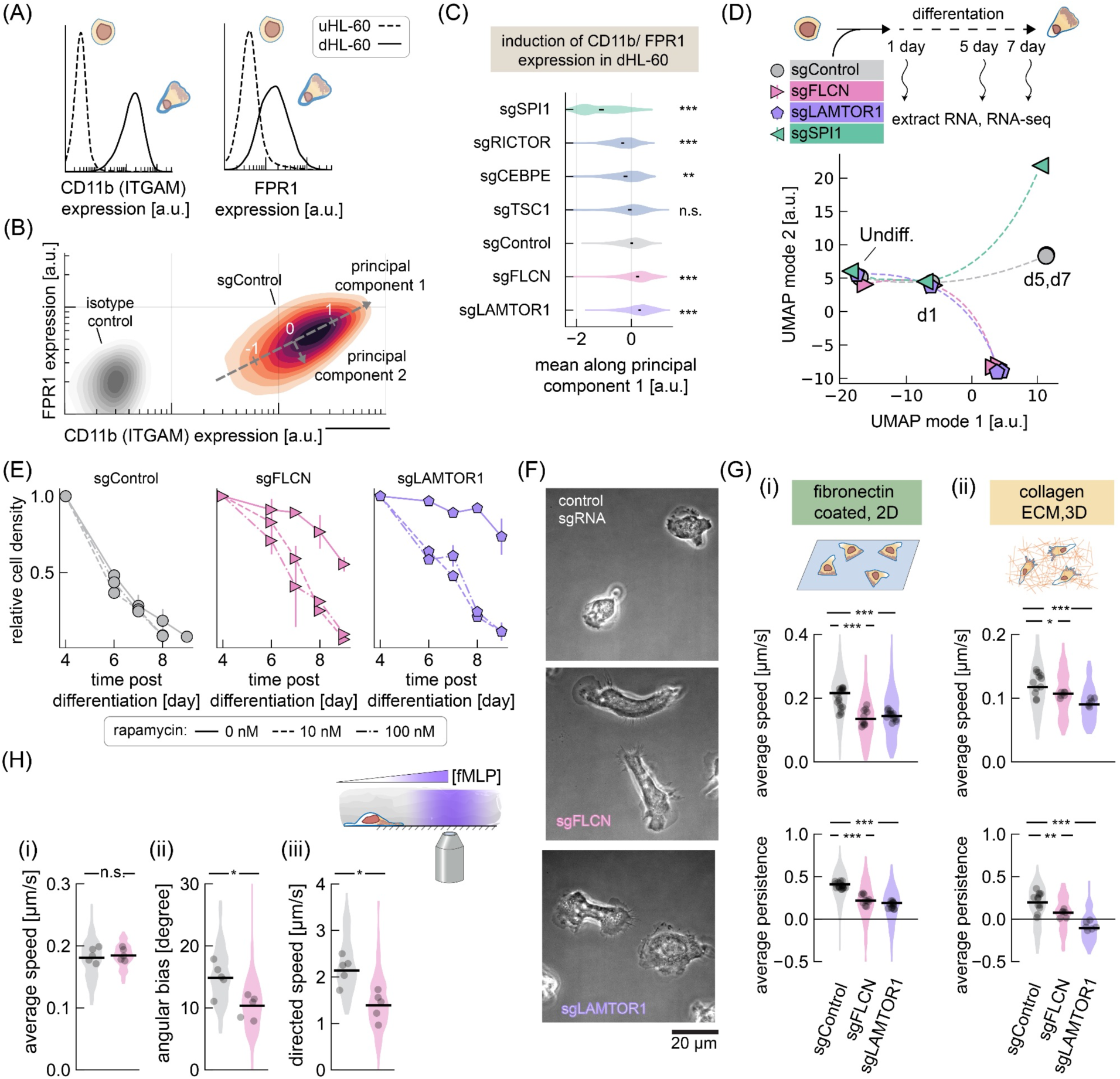
Knockdown FLCN and LAMTOR1 alters cell survival, with cells exhibiting increased adhesion, reduced migratory persistence, and poorer chemotactic sensitivity. (A) Flow cytometry immunofluorescence measurements of CD11b (ITGAM) and fMLP receptor (FPR1) cell surface expression in uHL-60 and dHL-60 cells. (B) The two-dimensional heatmap shows the induced expression of CD11b and FPR1 in dHL-60 cells. The axis associated with induction of these surface markers were identified by applying principal components analysis. Measurements using isotype control antibodies are shown in gray. (C) The first principal component identified in (B) was used to compare changes in expression induction in different gene knockdown lines. Black lines indicate the 99% confidence interval for the log expression mean along the first principle component, calculated by bootstrapping across single-cell flow cytometry measurements. (D) Transcriptional changes following knockdown of FLCN, LAMTOR1, and SPI1 were assayed by RNA-seq pre-differentiation (undiff.), 1-day, 5-day, and 7-day post-differentiation. Dimensionality reduction using UMAP was applied to transcription data and pseudo-plotted using a spline to show temporal trajectory. Individual data points represent an average across 6 RNA-seq replicate samples. (E) Rapamycin treatment targeting mTORC1 in dHL-60 CIRSPRi knockdown lines targeting FLCN, LAMTOR1, and a control sgRNA. Treatment of dHL-60 cells was begun on day 4 following the initiation of differentiation. Cells were either untreated (solid line), or treated with rapamycin at 10 nM rapamycin (dashed lines) and 100 nM rapamycin (dash-dot lines). Error bars represent standard deviation across replicates (N=3-4). (F) Phase microscopy of cell migration on fibronectin-coated coverslips with dHL-60 CIRSPRi knockdown lines targeting FLCN, LAMTOR1, and a control sgRNA. (G) Characterization of cell migration phenotypes following knockdown of FLCN and LAMTOR1. Speed was calculated by tracking cell nuclei during migration on fibronectin-coated coverslips (I) or in collagen ECM (ii). Persistence was inferred from the cell velocity data as described by Metzner et al. (see *Methods*). Individual data points represent average values for cells across a single field of view, with the shaded regions showing the distribution of all measurements. Measurements represent experiments performed over 2-3 days. (H) The acute chemotaxis response of dHL-60 cells was assayed by photo-uncaging fMLP during migration on BSA passivated coverslips, with cells confined by agarose. Average instantaneous speed (i), angular bias (ii), and the directed speed (projected speed along direction of fMLP gradient) (iii) are shown. Data points indicate average values across 5 different experiments (∼3,500 cells per cell line, per experiment), with the shaded regions showing the distribution across measurements .

These results supported the hypothesis that disruption of mTOR-related signaling might enhance neutrophil differentiation. In order to explore whether disruption of these genes might alter phosphorylation of known targets of mTORC1 and mTORC2 protein kinase activity, we performed Western blot assays to assess the phosphorylation status of the ribosomal S6 kinase and Akt, which are well-characterized targets of mTORC1 and mTORC2, respectively. Because the gene products of both FLCN and LAMTOR1 mediate mTORC1 kinase activity (Fig. 2C), we expected that knockdown of these genes would result in decreased phosphorylation of S6K. However, we found that phosphorylation of both S6K and Akt were largely unchanged in both our LAMTOR1 and FLCN knockdown lines (Fig. S3B-C), indicating that these perturbations did not substantially inhibit mTOR-related protein kinase activity.

It is known that mTORC1 can regulate the activity of transcription factors, including TFEB and TFE3, via their phosphorylation that is dependent on RagC/D-FLCN and mechanistically distinct from other well-characterized mTORC1 targets ^52, 53^. Notably, knockdown of TFEB and TFE3 resulted in a modest but significant decrease in sgRNA abundance in our differentiation screen, which contrasted with the positive enrichment of FLCN and LAMTOR genes (Fig. 2B). We, therefore, turned to whole-transcriptome sequencing (RNA-seq) to identify global transcriptional changes that might provide more insight into the changes in mTORC signaling. Focusing on the stable cell lines expressing sgRNAs targeting LAMTOR1, FLCN, and SPI1, we performed RNA-seq on uHL-60 cells, and dHL-60 cells at day 1, 5, and 7 following the initiation of differentiation. Using dimensionality reduction (UMAP ^54^) to take a broad look at the entire data set, we found similar changes in transcription in our LAMTOR1 and FLCN knockdown lines, distinct from both the line expressing our control sgRNA and the line expressing SPI1 sgRNA (Fig. 3D). For all cell lines, the transcriptional changes associated with differentiation following DMSO treatment appeared to stabilize in 5-day dHL-60, with little change observed in 7-day dHL-60 cells. Notably, the transcriptional profiles across our knockdown lines only diverged during differentiation, but were largely indistinguishable in undifferentiated cells. This is consistent with our screen results and importantly, shows that these genes are specifically involved in differentiation and neutrophil function.

To better understand why the FLCN and LAMTOR1 knockdown lines enjoyed a prolonged lifespan as healthy differentiated cells, we delved more deeply into the identities of the genes whose expression was altered relative to our control sgRNA line. Here, we found enrichment for genes associated with lysosomes, autophagy, and transcription of ribosomal genes (Fig. S3D-E). Importantly, these match the reported roles of TFEB and TFE3 as master regulators of lysosomal biogenesis and autophagy ^52^, further confirming that altered signaling is indeed along the RagC/D-FLCN axis of the mTORC1 signaling pathway. In addition, we found a substantial decrease in expression of the autophagy-activating kinases ULK1 and ULK2, and increased expression of the anti-apoptotic gene BCL2 (Fig. S3E). Interestingly, we also found significant changes in genes associated with degranulation, a key neutrophil immune function. More specifically, among gene products associated with primary granules (also known as azurophilic granules, mainly involved in phagosome formation), we found increased expression of myeloperoxidase and cathepsins, and a decrease in elastase expression. Within gene products associated with tertiary granules, which are exocytosed, we found a large decrease in the expression of matrix metallopeptidase 9 (MMP9).

Lastly, as an independent method to assay the functions of mTOR (*m*ammalian *T*arget *O*f *R*apamycin)-related signaling on HL-60 differentiation, we treated our stable cell lines with 10 nM and 100 nM rapamycin, dosages expected to abolish mTORC1 kinase activity ^55^. Rapamycin treatment of stable cell lines expressing sgRNAs targeting FLCN and LAMTOR1 resulted in a decrease in the survival of these cells following differentiation, restoring their survival characteristics to the lower levels associated with the sgRNA control cell line (Fig. 3E). This result strongly suggests that mTORC1 signaling is directly involved in cell survival after differentiation. In sum, these data help explain the observed pattern of enrichment of dHL-60 cells in our differentiation screen, where disruption of mTORC1-related genes alter (but do not eliminate) mTORC1 signaling, resulting in a reduction in apoptotic signals and extended neutrophil survival. This contrasts with rapamycin-treated cells, where a loss of mTOR activity in the FLCN and LAMTOR1 knockdown lines return viability back to their normal phenotype.

### Knockdown of folliculin and LAMTOR1 increase substrate adhesion but reduce migratory persistence and chemotaxis sensitivity

In our differentiation screen, knockdown of Ragulator, FLCN, or TSC1/2 all led to a similar enrichment of dHL-60 cells within the pooled CRISPRi library (Fig. 2C). Interestingly, however, only knockdown along the Ragulator-FLCN-Rag axis of mTORC1 signaling led to a perturbation across our cell migration assays (Fig. 2B, right panel). To better understand how knockdown of these genes influences migration and chemotaxis, we measured detailed cell motility characteristics for our stable knockdown lines targeting LAMTOR1 and FLCN as compared to control cell lines.

We began by looking at 2D cell migration on fibronectin-coated coverslips. Consistent with the elevated levels of CD11b in our LAMTOR1 and FLCN knockdown lines, these cells appeared more adherent. Specifically, in contrast to our control cell line expressing a non-targeting sgRNA, the LAMTOR1 and FLCN knockdown cells often lacked the normal front-back polarity, possibly due to inadequate contraction at the cell rear relative to their increased substrate adhesion (Fig. 3F). From our RNA-seq data above, these knockdown lines also exhibited a 2.5-fold increase in transcript expression of the integrin *α*_x_ (ITGAX), which like CD11b (integrin *α*_M_) pairs with integrin β_2_ and supports cell adhesion ^56–58^. Tracking cell nuclei as cells migrated on fibronectin-coated coverslips for 30 minutes, we found that the knockdown cells moved at about half the speed of our control sgRNA line, with an average speed of 0.13 µm/s as compared to 0.21 µm/s (Fig. 3G, i, top). We also estimated cellular persistence, employing a Bayesian inference algorithm based on a model for a heterogeneous random walk ^59^ to calculate a migratory persistence metric for each cell. In this model, a persistence of zero corresponds to a random walk. In line with the reduced front-back polarity, FLCN and LAMTOR1 knockdown lines showed a reduction in persistence (Fig. 3G, i, bottom) as compared to controls. In addition, we performed a series of assays to measure cell speed and persistence for spontaneous migration in 3D collagen gels. In the 3D assays, the LAMTOR1 and FLCN knockdown cells also exhibited a substantial loss of directional persistence, as well as a modest but statistically significant decrease in cell speed, moving about 10-25% slower than our control sgRNA line (Fig. 3G, ii).

We were also interested in whether the knockdown cells with altered mTORC1 signaling activity maintained their sensitivity to fMLP as a chemotactic agent, and explored this further in our FLCN knockdown line. Here we used photo-activation of caged fMLP to generate spatial gradients of the small chemotactic peptide ^30^, using a standard assay where migratory cells are sandwiched between a BSA-coated coverslip and an agarose overlay to minimize requirements for integrin-based adhesion during migration ^60^. While FLCN knockdown cells migrated with similar speeds as our control sgRNA line in this context (Fig. 3H,i), knockdown cells were much less responsive to fMLP. Across cells, we observed an average angular bias of about 10°, compared to 15° for our control sgRNA line (Fig. 3H, ii), which also resulted in a reduced directed speed (i.e. projected speed along the spatial fMLP gradient) (Fig. 3H, iii). Since fMLP receptor expression was higher in the FLCN knockdown cells (Fig. 3C, Fig. S3A), the defect in chemotaxis must lie downstream of receptor signaling.

In summary, in addition to the altered differentiation characteristics following disruption of Ragulator-FLCN-Rag signaling, these perturbations substantially alter adhesion and directed cell migration for dHL-60 cells.

### Migration through track-etch membranes is dominated by genes associated with cell adhesion

Thus far, we have established the utility of our whole-genome CRISPRi knockdown library in confirming known molecular mechanisms contributing to cell proliferation and differentiation, as well as identifying new and unexpected connections between differentiation and migratory behavior for dHL-60 cells. Next, we analyzed our several cell migration screens to compare the genes identified across the different migration contexts. One of the most notable observations was a strong correlation between the normalized log_2_ fold-change values for our chemokinesis and chemotaxis screens (Fig. 4A, left panel; ρ = 0.99), consistent with a proposition that there are no distinct molecular pathways that are specifically required for directional as opposed to random neutrophil migration in the context of serum stimulation. Among the knockdowns with the most detrimental effects on migration across these two assays were genes associated with inside-outside ɑ_M_β_2_ integrin signaling (Fig. 4B), highlighting the importance of adhesion as a key prerequisite to entry and migration through the pores of the track-etch membrane. Indeed, when we looked at migration of a stable knockdown line expression a sgRNA targeting ITGB2, cells exhibited a polarized morphology but were only loosely adherent during imaging of dHL-60 cells on a fibronectin-coated coverslip (Fig. 4C and Fig. S4A), a phenotype that is the opposite of the hyperadhesion exhibited by the FLCN and LAMTOR1 knockdowns described above. These results are broadly consistent with reports in other cell types that there is an optimum degree of cell-substrate adhesion for efficient migration, with either increasing or decreasing adhesion causing decreased cell speed ^61–64^.

**Figure 4:**
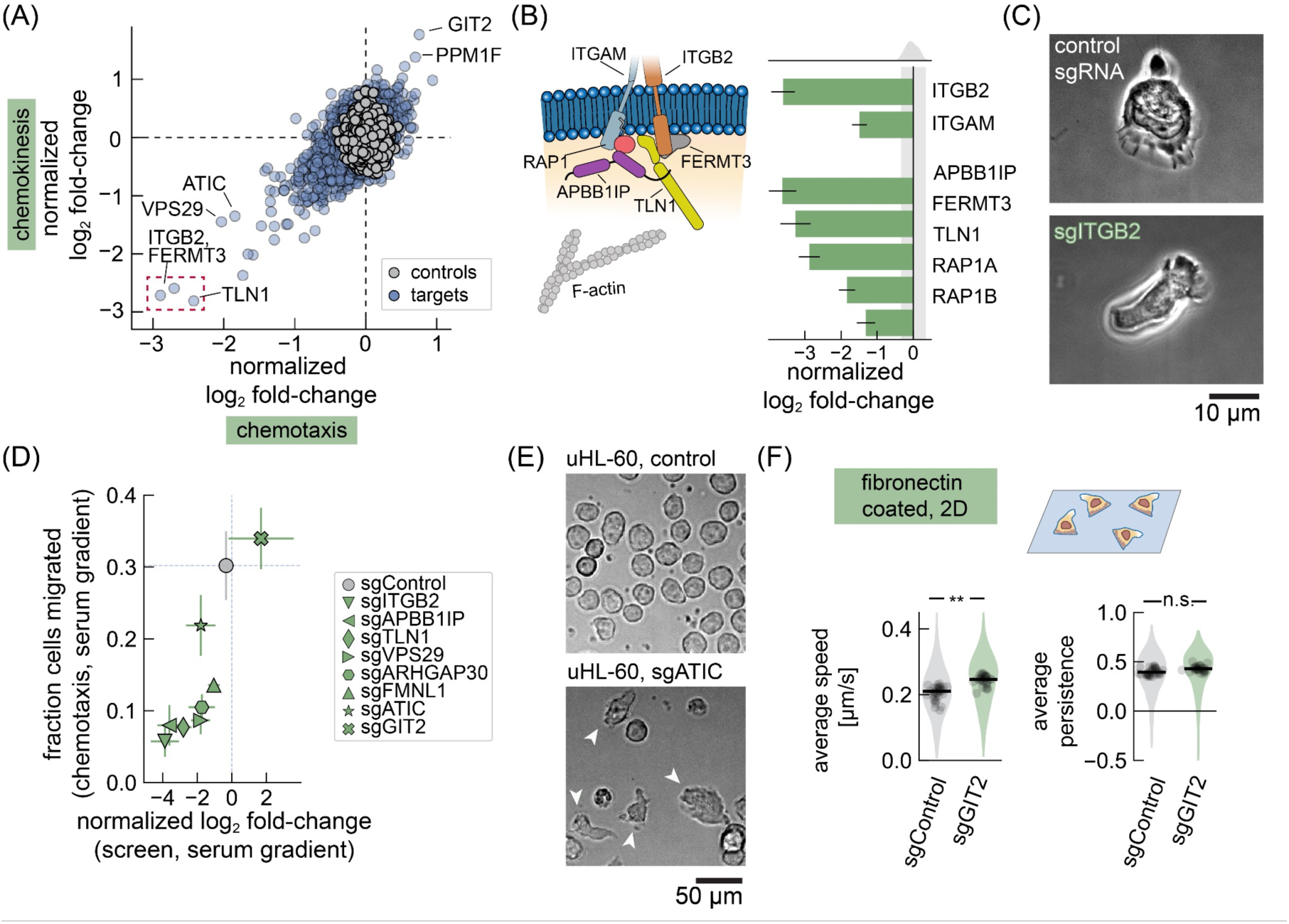
Cell migration CRISPRi screen identifies genes important for adhesion and migration on 2D surfaces. (A) Comparison of normalized log_2_ fold-changes across the pooled CRISPRi cell migration screens of chemotaxis and chemokinesis. Genes associated with inside-out *α*_M_*β*_2_ integrin signaling are among most significantly enriched across both assays, with ITGB2, FERMT3 and TLN1 genes identified. (B) Schematic of the key components of inside-out *α*_M_*β*_2_ integrin signaling and summary of their most significant sgRNA normalized log_2_ fold-changes values, averaged across all the chemotaxis and chemokinesis screens. Error bars represent standard error of the mean across individual screen experiments (N = 14). The gray shaded region shows the histogram of control sgRNAs. (C) Phase microscopy of cell migration on fibronectin-coated coverslips with dHL-60 CIRSPRi knockdown lines targeting ITGB2 and a control sgRNA. The elongated morphology of the ITGB2 knockdown line is because the cell is detached from the coverslip. Time courses are shown in Figure S4. (D) Comparison of chemotaxis normalized log_2_ fold-changes with measurements in individual sgRNA knockdown lines. Individual cell lines were exposed to a serum gradient by adding 10% hiFBS to the bottom-side of the track-etch membrane and allowing cells to migrate for two hours. The gray data point and dashed lines represent the values obtained for cells with a control sgRNA. Error bars represent standard error of the mean across replicates (N = 4 for individual transwell experiments using single sgRNAs). (E) Brightfield microscopy of uHL-60 CIRSPRi knockdown lines targeting ATIC and a control sgRNA. Control uHL-60 cells exhibit an expected round morphology, while sgRNA targeting of ATIC resulted in many cells that exhibited a migratory capability (white arrows). (F) Characterization of cell migration phenotypes following knockdown of GIT2. Speed was calculated by tracking cell nuclei during migration on fibronectin-coated coverslips. Persistence was inferred from the cell velocity data as described by Metzner et al. (see *Methods*). Individual data points represent average values for cells across a single field of view, with the shaded regions showing the distribution of all measurements. Measurements represent experiments performed over 3 days.

Given the correlation between our chemokinesis and chemotaxis screens, we wanted to relate the measured log_2_ fold-change values with a more biologically relevant parameter associated with cell migration. One of the challenges with using fold-changes in sgRNA abundance to identify significant biological perturbations is that it provides little initial insight into the magnitude of biological effect. To this end, we constructed stable cells lines expressing sgRNAs that spanned the range of observed log_2_ fold-change values (ITGB2, APBB1IP, TLN1, VPS29, ARHGAP30, FMNL1, ATIC, GIT2) and performed individual transwell assays with 10% hiFBS added to the bottom reservoir, similar to the conditions for our larger-scale chemotaxis screen. Quantifying the fraction of cells that migrated through the track-etch membrane after two hours, we found a strong correlation with the log_2_ fold-change values from our chemotaxis screen (Fig. 4D, ρ = 0.87). More specifically, the strongest perturbation, targeting knockdown of ITGB2, led to only 6% of the cells in the bottom reservoir versus 30% of the cells with our control cell line. Knockdown of GIT2 showed the largest positive increase, with 34% of the cells collected in the bottom reservoir.

We also noted that during our construction of a stable cell line with a sgRNA targeting ATIC, a substantial number of polarized migratory cells were present even prior to treatment with DMSO, a striking phenotype that is not observed in unperturbed uHL-60 cells (Fig. 4E and Fig. S4B). RNA transcriptome analysis showed that these cells had a transcriptional profile more similar to that of dHL-60 cells with a control sgRNA (Fig. S4C) than to control uHL-60 cells, suggesting that ATIC knockdown was causing spontaneous differentiation. ATIC codes for an enzyme that acts on the adenosine monophosphate analog AICAR, an intermediate in the generation of inosine monophosphate in the purine biosynthesis pathway ^65^. AICAR is capable of stimulating AMP-dependent protein kinase (AMPK) activity ^66, 67^ and one possibility is that ATIC knockdown is changing the basal concentration of AICAR which may alter the metabolic and energy state in the uHL-60 cells and drive differentiation.

Since most gene knockdowns decreased the fraction of cells migrating through the track-etch membrane, we were particularly intrigued by the smaller subset of genes that exhibited a positive log_2_ fold-change in our chemotaxis and chemokinesis screens; that is, those whose knockdown enhanced cell migration. Among the most positively enriched genes was GIT2 (Fig. 4A), which encodes a protein that binds to the p21-activated kinase-interacting guanine nucleotide exchange factors α-PIX (ARHGEF6) and β-PIX (ARHGEF7). α-PIX and β-PIX both enhance the activity of the Rho GTPases Cdc42 and Rac1 ^68, 69^ that act as master regulators to enhance actin assembly at the leading edge of motile cells ^70^. In mouse neutrophils, loss of GIT2 was previously shown to lead to impaired chemotactic sensing and enhanced superoxide production ^71^. Notably, knockdown of α-PIX and another α-PIX binding partner, the Ser/Thr protein phosphatase PPM1F, also exhibited positive enrichment in our migration screens (Fig. 4A). To better understand how knockdown of GIT2 influenced cell migration, we directly examined the motility behavior of our stable cell line with a sgRNA targeting GIT2. Analyzing cell tracks as cells migrated on fibronectin-coated coverslips, we find that GIT2 knockdown cells migrated with an average speed of 0.25 µm/s, or about 20% faster than our control cell line (Fig. 4C, left). Migration otherwise appeared similar to control cells, exhibiting similar migratory persistence (Fig. 4C, right).

### CRISPRi screen identifies genes important for 3D amoeboid cell migration

We next turned to the results from our screen of 3D amoeboid migration. In comparison with our track-etch based chemokinesis screen, many of the genes associated with adhesion did not exhibit strong phenotypes in the 3D migration assay (Fig. 5A), consistent with expectations for an integrin-independent mode of cell migration for cells embedded in fibrous ECM ^72^.

**Figure 5:**
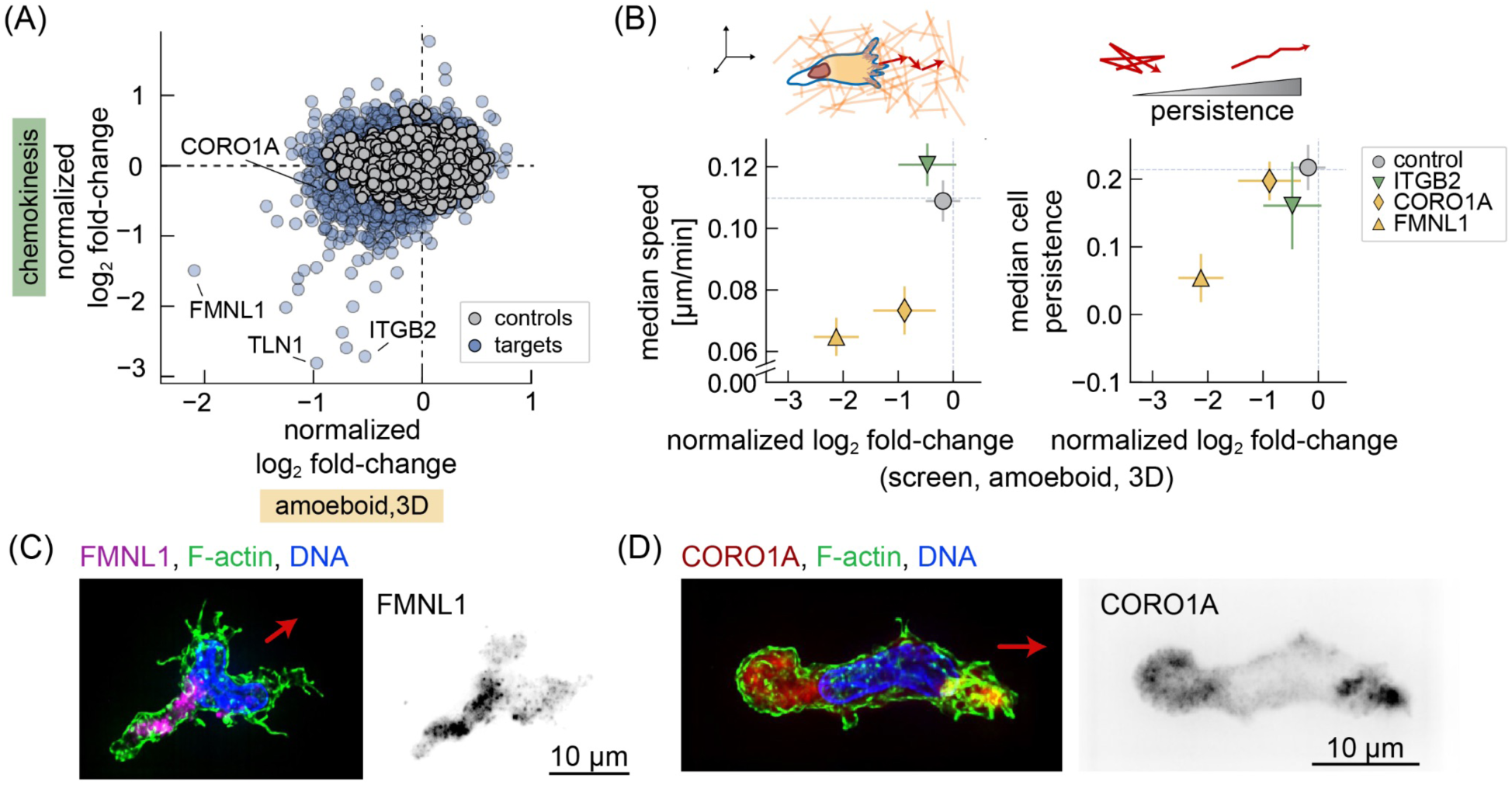
Cell migration CRISPRi screen identifies genes important for amoeboid, 3D migration. (A) Comparison of normalized log_2_ fold-changes across the pooled CRISPRi cell migration screens of amoeboid, 3D migration and chemokinesis. (B) Comparison of amoeboid 3D normalized log_2_ fold-changes with measurements of cell speed and migratory persistence via single-cell nuclei tracking. Cells from individual sgRNA knockdown lines were tracked during migration in collagen for 60 minutes (1 minute frame rate). The median cell speed (left) and inferred cell persistence (right, see methods) are plotted against their measured normalized log_2_ fold-change from the pooled screen. Error bars represent standard error of the mean across replicate experiments (N = 6 for screens and N= 4 for experiments using single sgRNA cell lines). (C) & (D) show immunofluorescence localization of FMNL1 and CORO1A in amoeboid-migrating cells in collagen. F-actin was labeled by phalloidin, while DNA was stained by DAPI. Images are maximum projections; right panels show grayscale localization of formin-like 1 (C) and coronin 1A (D). Red arrows indicate the direction of cell migration.

Interestingly, knockdown of talin 1 (TLN1), which mediates the linkage between integrins and the actin cytoskeleton, still produced a significant phenotype in the 3D amoeboid screen. This suggests additional roles for talin 1 beyond the interaction between the actin cytoskeleton and ɑ_M_β_2_ integrins in dHL-60 cells.

We chose to further characterize two actin regulatory proteins identified in the 3D amoeboid screen, formin-like 1 (FMNL1) and coronin 1A (CORO1A). These proteins have been implicated in cell migration ^73–75^, and FMNL1 was also identified as a significant hit in our chemotaxis/ chemokinesis screens above, but their role during 3D migration remains less well-characterized. We generated stable cell lines with sgRNA targeting these genes to quantify migratory speed and persistence as cells migrated in 3D collagen gels. As expected, knockdown of ITGB2 showed no effect on speed or persistence in 3D, but migration speed was decreased in FMNL1 and CORO1A knockdowns, consistent with the results of the 3D screen (Fig. 5D, left panel, ρ = 0.83). For both knockdown lines, cells migrated with an average speed of approximately 0.07 µm/s, roughly half as fast as our control sgRNA or ITGB2 knockdown lines, which have average speeds of 0.11 - 0.12 µm/s.

While knockdown of either FMNL1 or CORO1A resulted in reduced speed, only FMNL1 knockdown showed a significant reduction in migratory persistence (Fig. 5B, right panel). This may reflect different roles during 3D migration. We used immunofluorescence to determine the localization of these proteins in wild-type cells. In contrast to the expected localization of well-characterized formins like mDia1/2 ^76^, we find FMNL1 rear-localized and often directly behind the nucleus (Fig. 5C). This appears consistent with recent work in T-cells ^77^, who found similar localization and hypothesized that formin-like 1 may support actin polymerization to aid in squeezing the nucleus through tight endothelial barriers. In the 3D context considered here, formin-like 1 may be important for migratory persistence by supporting movement through the collagen gel. In contrast to formin-like 1, we find coronin 1A predominantly colocalized with the lamellipodial filamentous actin structures at the cell front, though more diffuse protein localization was also observed at the cell rear (Fig. 5D). Coronin 1A localization to the lamellipodial projections appear consistent with the characterization of cells migrating on a 2D surface ^75, 78, 79^, while the rear-localized protein may relate to a role in actin turnover and disassembly ^73^. Here, the lack of change in migratory persistence following CORO1A knockdown may relate to a more general disruption of actin cytoskeleton dynamics, rather than alterations to how cells move through the collagen ECM, though further work will be needed to clarify this point.

### Differential sensitivity to protein trafficking machinery and integrin expression is observed across cell migration assays

One gene not obviously associated with cytoskeletal function that was among those whose knockdown was most disruptive to migration across all screens was VPS29, a component of the retromer and retriever complexes (Fig 4A and Fig. 5A). These complexes recycle transmembrane proteins from endosomes back to the trans-Golgi network and the plasma membrane, respectively ^80, 81^. Exploring these complexes further, additional subunits for both the retromer and retriever complexes were identified as significant hits in our migration screens (Fig. 6A, left). Among other proteins involved in protein trafficking, we also identified components of the HOPS and CORVET complexes (Fig. 6A, right), which are specifically involved in endosomal-lysosomal protein trafficking^82^ and may also influence mTORC1 signaling^52^.

**Figure 6:**
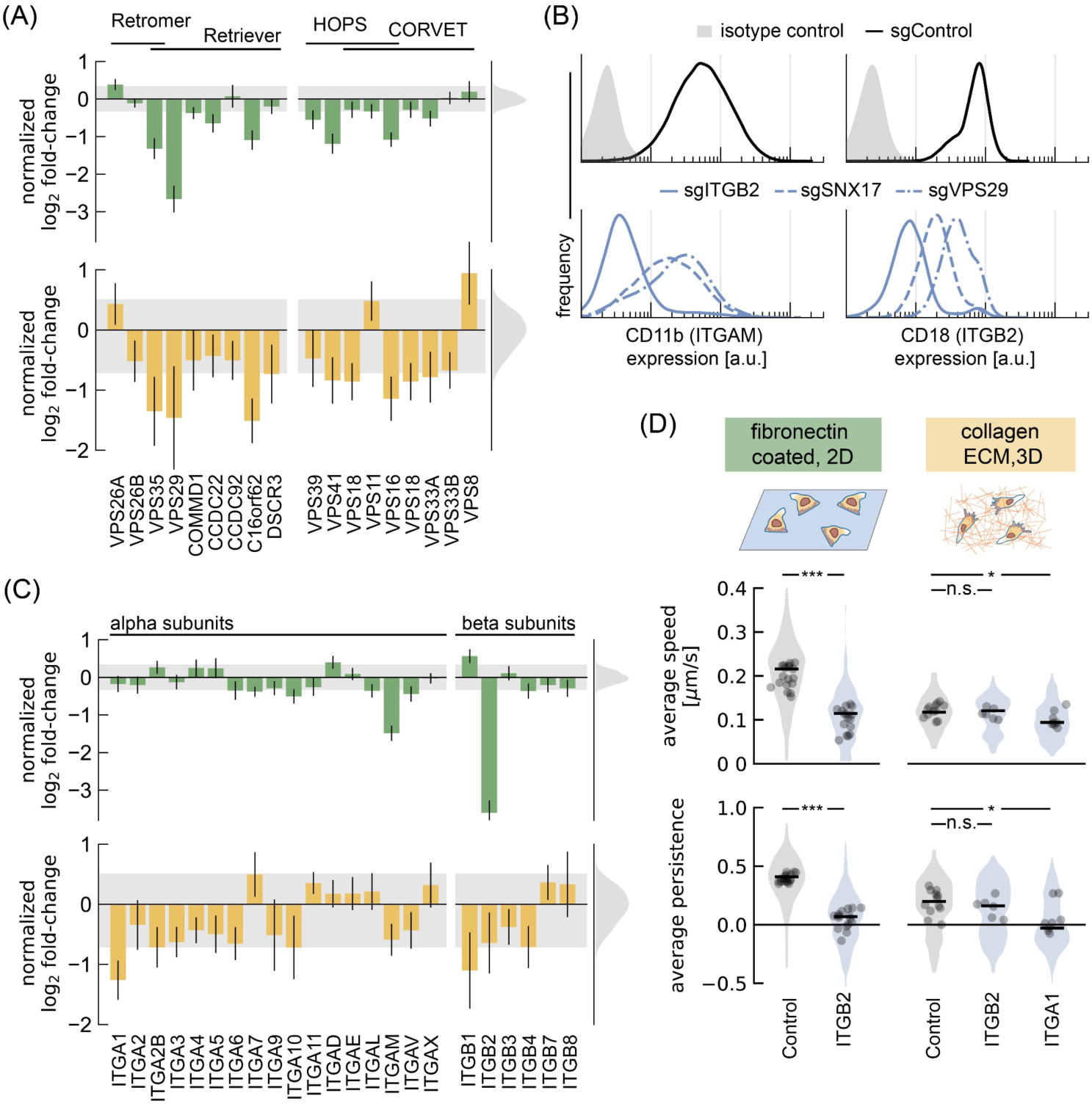
2D and 3D migration show context specific sensitivities to integrin expression and recycling. (A) Summary of normalized log_2_ fold-changes for protein trafficking genes in the chemotaxis and chemokinesis screens (green), and the amoeboid screen (yellow). Shared gene products for the protein complexes (retromer/ retriever and HOPS/ CORVET) are indicated by a solid black line. Error bars represent standard error of the mean across screen replicates, where the most significant sgRNA perturbation from each gene. The histograms and shaded region identify the distribution of the control sgRNAs. (B) Immunofluorescence flow cytometry of CD11b (ITGAM gene; left column) and CD18 (ITGB2 gene; right column). Histograms show surface distribution in control sgRNA (black, solid), ITGB2 (blue, solid), SNX17 (blue, dashed), VPS29 (blue, dash-dot). Shaded histograms indicate cellular autofluorescence from an non-targeting isotype control antibody. (C) Summary of normalized log_2_ fold-changes for integrin genes in the chemotaxis and chemokinesis screens (green), and the amoeboid screen (yellow). Integrin genes whose transcription is not detected in HL-60 cells^86^ were excluded. Error bars represent standard error of the mean across screen replicates, where the most significant sgRNA perturbation from each gene.. The histogram and shaded region identify the distribution of the control sgRNAs. (D) Characterization of cell migration phenotypes in integrin knockdown lines. Speed was calculated by tracking cell nuclei during migration on fibronectin-coated coverslips or in collagen ECM. Persistence was inferred from the cell velocity data as described by Metzner et al. (see *Methods*). Individual data points represent average values for cells across a single field of view, with the shaded regions showing the distribution of all measurements. Measurements represent experiments performed over 2-3 days.

Given the importance of cell-substrate adhesion in cell migration, and particularly for our chemotaxis/chemokinesis screens, we considered the hypothesis that perturbations affecting protein trafficking in the membrane recycling pathway might alter integrin recycling and degradation ^83^. Further supporting this, sorting nexin 17 (SNX17), which bind to ꞵ integrins in conjunction with the retriever complex to recycle integrins back to the plasma membrane ^84^, was also identified in our chemotaxis and chemokinesis screens. To test whether integrin expression was altered when genes associated with membrane recycling were knocked down, we generated additional cell lines with sgRNAs targeting VPS29 and SNX17 and measured their cell surface expression of ɑ_M_β_2_ integrins (CD11b and CD18, for integrin ɑ_M_ and β_2_, respectively) using flow cytometry. As a positive control, we found that the ITGB2 knockdown cell line showed a near complete loss of β_2_ integrin expression, as expected (Fig. 6B, right). Only assembled heterodimer ɑβ integrin pairs are expected to be stably localized at the cell surface ^85^, and consistent with this, these cells also exhibit a near-complete loss of integrin ɑ_M_ (Fig. 6B, left). Examining our cell lines with sgRNA targeting VPS29 and SNX17, we found a moderate drop in ɑ_M_β_2_ integrin expression relative to our control sgRNA (Fig. 6B), with measurable decreases in the expression levels of both subunits. These findings show that the migration defects associated with perturbations to the membrane recycling pathway may be due, at least in part, to disruptions in integrin surface presentation.

Interestingly, although 3D amoeboid migration was insensitive to knockdown of ɑ_M_β_2_ integrin (Fig. 5B), several genes among the protein trafficking complexes, including VPS29, still disrupted cell migration when knocked down in our ECM screen (Fig. 6A). Although disruption of these protein complexes may have other pervasive effects beyond altering integrin expression, we wanted to explore integrins more comprehensively across our cell migration screens by comparing the expression of all integrin subunits that exhibit measurable transcripts in these cells ^86^. Notably, in our 3D amoeboid screen, sgRNA targeting ITGA1 led to a significant perturbation to cell migration (Fig. 6C). Integrin heterodimers containing the integrin ɑ_1_ subunit are known to bind collagen ^87^, which was used as the main ECM component in this screen. To further validate this perturbation, we constructed an individual cell line with sgRNA targeting ITGA1 and tracked its migration in 3D. In contrast to ITGB2 knockdown, which only affected 2D migration, our line targeting ITGA1 showed a modest, but significant reduction in cell speed and a large drop in cellular persistence for 3D migration (Fig. 6D). In summary, we find that protein trafficking, and related alterations in integrin cell-surface expression, can alter behavior across both 2D and 3D cell migration.

### CRISPRi screens of cell migration provide a rich resource for studying rapidly migrating cells

Beyond the genes related to adhesion, protein trafficking, and those associated with the actin cytoskeleton noted above, our screen identified many other genes that may have either direct or indirect roles in cell migration (listed in Supplementary Data Tables 3, 4 and 5). For example, a variety of genes were identified whose proteins localize to the endoplasmic reticulum and Golgi apparatus. Further investigation will be needed to better understand their specific significance to cell migration. To explore our migration data sets more broadly, we also applied pathway enrichment analysis to the results of our cell migration screens (Fig. 7A). Combining this with our exploration of the data thus far, in Fig. 7B we provide a summary of genes identified across our cell migration assays.

**Figure 7:**
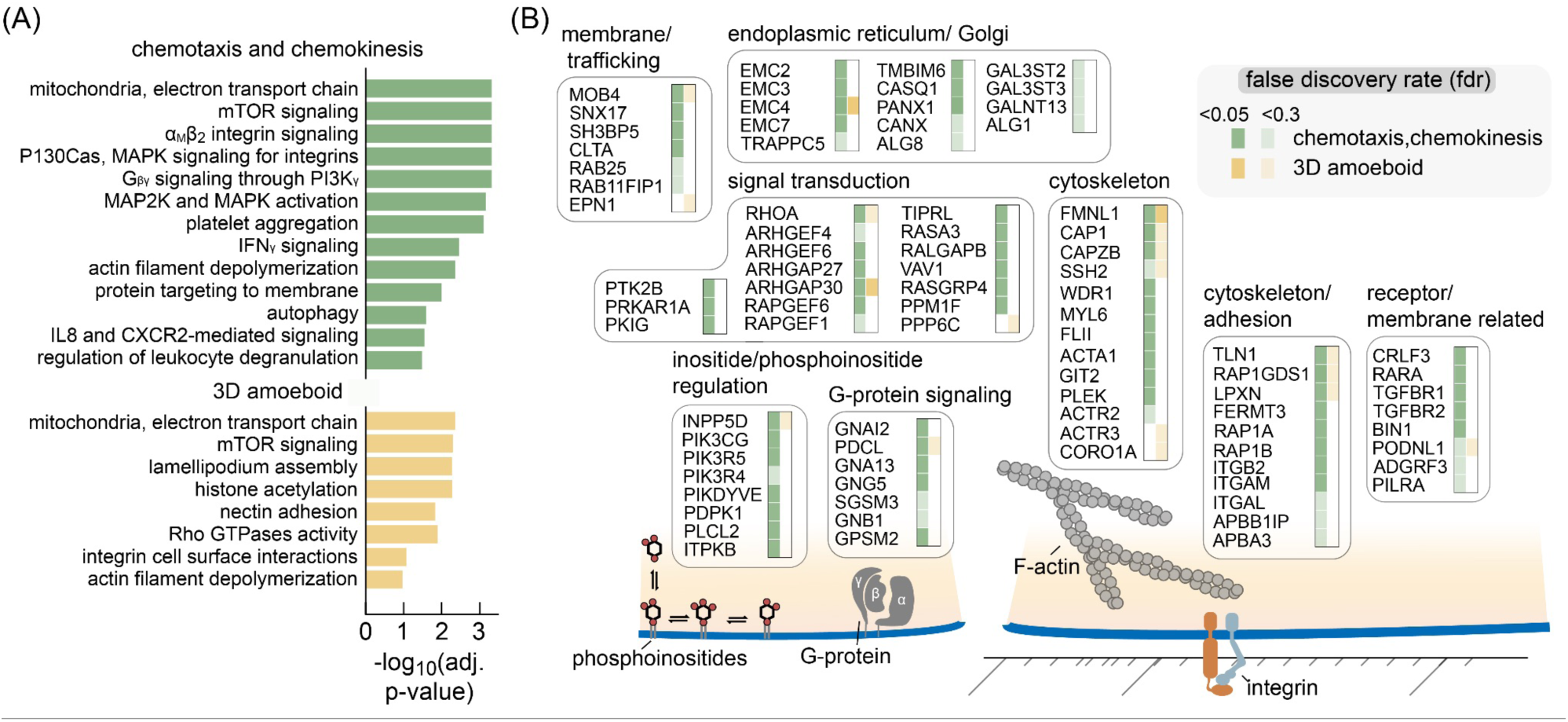
Summary of pathways and genes identified across cell migration CRISPRi screens. (A) Pathways enriched in cell migration screens. Since the majority of gene knockdowns lead to poorer migratory phenotypes (i.e. negative log_2_ fold-changes), disruption of the noted pathways are associated with poorer migration. Due to the correlation across the chemotaxis and chemokinesis screens, their data were combined in this analysis (green). Pathways enriched in the amoeboid, 3D screen are shown in yellow. (B) Summary of the genes identified across the cell migration screens. Genes were identified from the collated chemotaxis/ chemokinesis screens (green) and 3D amoeboid screen (yellow), with the shading intensity indicating the false discovery rate threshold that each gene fell into (<0.05 or <0.3). Empty column entries (i.e. white) mean that the gene was not significant. Genes associated with transcription, translation, and gene regulation were excluded. In addition, genes that perturbed the processes of either proliferation or differentiation, with absolute log_2_ fold-changes values larger than 0.7, were excluded.

Although we have focused on a number of genes that appeared specific to the different migratory contexts (e.g. integrin ɑ_M_β_2_ is much more important for migration across a track-etch membrane than for migration in a 3D collagen matrix), a variety of genes were identified across all three cell migration screens, with many appearing to disrupt key steps known to be associated with actomyosin based migration. For example, the ꞵ subunit of the filamentous-actin capping protein CapZ (CAPZB), which is known to cap actin filaments at their barbed ends and prevents both filament elongation and filament depolymerization ^88^, had a measurable effect in all migration screens. The adenylyl cyclase-associated protein 1 (CAP1) was also identified in all three migration screens; this regulatory protein facilitates cofilin-driven actin filament turnover and may also interact with talin 1 ^89^. Although their phenotypes in the screens were quantitatively modest, several subunits of the nucleator of branched-actin polymerization, ARP2/3 were also identified. Specifically, ACTR2 was identified in our chemotaxis track-etch-based screen, while ACTR3 showed up in our amoeboid 3D screen. With respect to myosin contraction at the cell rear, knockdown of RhoA, a key regulator of myosin activity, significantly perturbed migration across all three screens. Among modulators of Rho GTPases, knockdown of ARHGAP30 was among the most significant perturbations and has been reported to negatively regulate activity of RhoA and Rac1 through enhancing GTP hydrolysis ^90, 91^.

## Discussion

The rapid migratory characteristics of neutrophils and their early role in our innate immune response to infection or wounds make them an important cell type to consider in the context of cell migration. We have demonstrated the use of pooled genome-scale gene perturbations using CRISPRi gene knockdown to study cell migration in human dHL-60 neutrophils. This was made possible through the development of scalable assays that effectively separate cells based on their migratory capabilities. This included the scaled-up application of a transwell assay through the use of track-etch membranes, where we compared chemotaxis and chemokinesis in the context of serum-stimulated cell migration. In the context of 3D amoeboid cell migration, which remains much less studied ^92^, we devised a multilayer collagen/fibrin gel that enabled isolation of the most migratory cells at scale. In addition, we performed CRISPRi screens of proliferation and neutrophil differentiation that provided important additional dimensions to distinguish biological roles across the processes of cellular growth, differentiation, and cell migration.

While the results from our screen of differentiation may reflect genetic sensitivities of the HL-60 leukemia cell line and our differentiation protocol, we were also encouraged by several results that showed our perturbations were also relevant to primary neutrophil differentiation, including the identification of genes encoding known master regulators for neutrophil differentiation such as CEBPA, CEBPE, and SPI1. Beyond regulatory targets, we found a strong enrichment for sgRNAs targeting genes involved in metabolism and oxidative phosphorylation. Intriguingly, we found substantial overlap of enriched genes when comparing our results with those from a differentiation screen for exit from pluripotency in mouse embryonic stem cells 42,43. Pathways regulating stem cell renewal and differentiation share substantial overlap with stress-response pathways ^93^ and our observation suggests these processes continue to be important as mammalian cells change their proliferative status during myeloid differentiation. In the context of myeloid cells, terminal cell differentiation would provide a useful strategy by the host to minimize the propagation of damaged or cancerous cells and our results appear particularly relevant to differentiation therapy as a treatment for patients with acute promyelocytic leukemia ^34^. Here it is worth noting that DMSO, which was used to induce differentiation into dHL-60 cells in our studies, can alter membrane permeability and impair mitochondrial function ^94, 95^. While the mechanism behind DMSO-mediated differentiation remains poorly understood ^15^, it may be that DMSO acts as a chemical insult that drives cell differentiation by impairing cellular homeostasis. Mitochondria, in particular, are hubs of metabolic signaling that produce molecules that modulate cellular function, gene expression, and can alter differentiation state ^96, 97^. Furthermore, the observation that knockdown of ATIC, encoding an enzyme required for purine biosynthesis, led to spontaneous differentiation of proliferating HL-60 cells into a migratory phenotype also appears consistent with the hypothesis that differentiation is particularly sensitive to disruption of metabolic processes. For ATIC knockdown specifically, this may be through alterations in energy homeostasis through altered regulation of AMP-dependent protein kinase (AMPK) activity ^66, 67^.

We found neutrophil differentiation and migration to be particularly sensitive to knockdown of genes associated with mTORC1 signaling. As a key regulator of cellular growth and metabolism, mTORC1 integrates many upstream signals that alter its kinase activity toward a host of downstream targets. Dysfunctional mTORC1 signaling is observed across many human diseases ^98^ and recent work has begun to uncover its role in our innate immune system ^99, 100^. In macrophages, knockout of genes involved in mTORC1 signaling substantially altered the phagocytosis efficiency toward a variety of targets ^11, 14^, possibly through alterations in the activation status of these cells ^100^. In our work, knockdown of genes involved in regulating mTORC1 activity led to an enrichment of differentiated dHL-60 cells. One possible explanation is that these cells proliferate more than control sgRNA cells during early differentiation, leading to an enrichment in the CRISPRi pool of dHL-60 cells, as could be the case with TSC1/TSC2 where gene knockdown is expected to constitutively activate mTORC1. Notably, knockdown of genes along the Ragulator-FLCN-Rag axis of mTORC1 signaling (i.e. LAMTOR1 and FLCN) resulted in cells that were also longer-lived and with substantially altered migratory behavior, leading to their identification in our screens of cell migration.

Western blot, RNA-seq analysis, and pharmacological treatment with rapamycin show that mTORC1 is still active following knockdown of LAMTOR1 and FLCN in HL-60 cells, albeit with altered target activity. Specifically, the phosphorylation state of the canonical mTORC1 target S6K was unchanged and instead we found substantial transcriptional changes associated with processes of lysosome biogenesis, autophagy, and cellular degranulation. This appears due to non-canonical mechanisms of mTORC1 substrate activation that have only recently begun to be elucidated ^52, 53, 101^. Here, signaling via FLCN-RagC/D played a dominant role in the regulation of catabolic processes, likely altering the activity of mTORC1 toward transcription factors like TFEB and TFE3 that are master regulators of lysosomal biogenesis and autophagy. In the context of neutrophils, our characterization of cell behavior after differentiation in individual knockdown lines of LAMTOR1 and FLCN showed major changes in the neutrophil lifetime, increased substrate adhesion, and poorer migratory persistence.

Our observations in the reduced chemotactic sensitivity and major transcriptional changes for genes involved in cellular degranulation following knockdown of LAMTOR1 and FLCN also suggest additional changes in downstream effector capabilities of these cells that will warrant further investigation. Indeed, emphasizing the relevance of these results from work in mice, inhibition of autophagy in neutrophils led to a reduction in degranulation in vivo and altered the neutrophil response in models of both inflammation and autoimmunity ^102^. mTORC1 signaling and autophagy is intimately tied to metabolism and energy production. With regard to neutrophil effector function, mitochondrial reactive oxygen species also influence the extent of NETosis induction, a dramatic event where neutrophils eject their DNA to entangle and kill invading pathogens ^103^. Metabolic defects also appear to contribute to neutrophil-associated diseases like chronic obstructive pulmonary disease (COPD) ^104^, while in some tumors, there is evidence for metabolically adapted, oxidative neutrophils that help maintain local immune suppression ^105^. These examples highlight the importance in understanding how alterations in metabolic signaling and processes like oxidative phosphorylation might further alter neutrophil behavior.

Our use of multiple cell migration screens allowed us to distinguish the potential roles of different genetic perturbations on migration in specific contexts, where we used migration across track-etch membranes to mimic extravasation of neutrophils as they initially exit the bloodstream, and used migration through 3D collagen and fibrin gels to mimic migration through solid tissues. Perhaps most striking was the clear importance of cell-substrate adhesion in our chemotaxis and chemokinesis CRISPRi screens using track-etch membranes. In particular, genes associated with inside-out *α*_M_*β*_2_ integrin signaling that are important for integrin activation and adhesion ^106, 107^ dramatically reduced migration success. These results are especially relevant to the family of leukocyte adhesion deficiency (LAD) disorders, where neutrophils are unable to effectively extravasate into tissue and mount an immune response following infection^108^. Notably, both ITGB2 and FERMT3, two of the strongest hits in our chemotaxis/chemokinesis screens, are also single-gene defects causing the LAD1 and LAD3 subtypes of this disorder, respectively ^109, 110^. The majority of these adhesion genes were not found to be important for 3D amoeboid migration, which is consistent with prior work showing that this type of cell migration is largely integrin-independent ^72, 111^. However, our 3D screen did allow us to identify a specific role for integrin ɑ_1_ (ITGA1), where we found that knockdown led to cells with poorer persistence when monitoring single cells migrating in a 3D collagen matrix. Overall, the results from our 3D amoeboid migration screen were notably distinct from the track-etch membrane screen results, highlighting the value in using multiple assays to probe cellular function in different, but related contexts. Knockdown of the formin gene, FMNL1, in particular, led to cells whose speed and migratory persistence were substantially lower than control cells. Given the rear-localization of FMNL1 and potential role for nucleus positioning ^77^, it will be interesting to explore how alterations in rear cell contractility or ECM composition and density alter migration characteristics in future work.

We observed a strong, indeed a near-perfect, correlation between the log_2_ fold-change values between chemotaxis and chemokinesis track-etch membrane experiments. As a first attempt to compare these two processes genome-wide, the lack of any notable enrichment for any genes as showing distinct results between the two assays suggests that there are no major genetic differences or distinct molecular pathways required for directional as opposed to random neutrophil migration in the context of serum stimulation. This was true even though our chemotaxis assay resulted in substantially more cells migrating to the bottom reservoir than the chemokinesis assay over equivalent time frames. This result speaks to the ability of neutrophils, as well as more disparate motile cell types like fish keratocytes ^112^, to spontaneously polarize their migratory machinery in the absence of any directional cue. This polarization is governed through the generation of well-documented front and rear signaling modules that are maintained through feedback loops ^113–115^. Chemotaxis, at least in the context of serum stimulation as explored here, likely acts to more efficiently guide the movement of cells through the pores of the track-etch membrane, but our results indicate that the underlying molecular mechanisms of directed and spontaneous cell motility are essentially identical. In future work it will be interesting to see how these results compare with more specific chemoattractants ^116^ or in the context of self-generated gradients ^117^.

In summary, our data provides a valuable resource for future study of proliferation, differentiation, and context-dependent cell migration of rapidly migrating neutrophils. Further experimental adjustments or the use of orthogonal modes of genetic perturbation such as gene activation ^9^ may provide additional insights into cell migration. For example, for our 3D amoeboid screen, changes could be made to the ECM composition and density ^92^ or spatial gradients could be implemented ^118^. These alternative experimental paradigms could be used to yield new insights into other modes of cell migration like durotaxis (gradients in ECM rigidity)^119^, haptotaxis (gradients in substrate composition)^120^, or galvanotaxis (directional response to electrical cues)^122^.

## Materials and Methods

### Cell Culture and Neutrophil Differentiation

Undifferentiated HL-60 cells (uHL-60) were cultured and differentiated into neutrophil-like cells as previously described ^19, 123^. Briefly, cells were maintained at 37°C and 5% CO2, cultured in RPMI 1640 medium containing L-glutamine and 25 mM HEPES 1640 (Gibco #22400089) supplemented with 10% heat-inactivated fetal bovine serum (hiFBS) (Gemini Bio Products #900–108), and 100 U/mL penicillin, 10 μg/mL streptomycin, and 0.25 μg/mL Amphotericin B (Gibco #15240). Differentiated HL-60 cells (dHL-60) were generated by incubating cells in media containing 1.57% Dimethyl Sulfoxide (DMSO, Sigma, #D2650). Here, confluent cells (approx. 1 - 1.5 x 10^6^ cells/mL) were diluted by adding two volumes of additional media and DMSO. The culture media was replenished with fresh media, including DMSO, three days following the initiation of differentiation. Except for where noted otherwise, dHL-60 cells were used in cell migration assays five days following the initiation of differentiation. Experiments involving rapamycin treatment used rapamycin (Thermo Scientific Chemicals #AAJ62473MF).

### CRISPRi pooled library construction

All genomic integrations involved lentiviral transduction. uHL-60 cells expressing dCas9-KRAB were first generated and this cell line was used for all subsequent work (sgRNA genome-wide libraries and individual sgRNA targeting cell lines).

The dCas9-KRAB construct was based on a construct originally gifted by Dr. Marco Jost and Dr. Jonathan Weissman, but modified to include blasticidin resistance (pHR-UCOE-Ef1a-dCas9-HA-2xNLS-XTEN80-KRAB-P2A-Bls).

The sgRNA library was previously reported in Sanson et al. 2018 (Dolcetto CRISPRi library set A, Addgene #92385). This library contains 57,050 sgRNA, with 3 sgRNA per gene target and 500 non-targeting control sgRNA. Individual sgRNA constructs used for single gene knockdown lines were generated based on sgRNAs from this library, with sgRNA sequences integrated into base vector pXPR_050 (Addgene #96925) as previously described^9^.

For large scale lentivirus production (dCas9 construct or pooled sgRNA library), 15 μg transfer plasmid, 18.5 μg psPAX2 (Addgene #12260), and 1.85 μg pMD2.G (Addgene #12259) were diluted in 3.5 ml Opti-MEM I reduced-serum media (Gibco #31985070) and then combined with 109 μL TransIT-Lenti Transfection Reagent (Mirus, MIR6600). Following a 10 minute incubation, this mixture was added dropwise to confluent HEK-293T cells (ATCC, CRL-3216) in a T175 flask containing 35 mL DMEM media (Gibco #11965-092) and supplemented with 1mM sodium pyruvate (Gibco #11360-070). Lentivirus was recovered by collecting media 48 hr later, with centrifugation at 500 g for 10 minutes to remove any residual cells and debris. For our dCas9-KRAB construct, we additionally concentrated the lentivirus approximately 60-fold using Lenti-X Concentrator (Takara Bio Inc., #631231).

For small scale lentivirus production (individual sgRNAs), lentivirus was prepared in 6-well tissue culture plates. Here 1 μg sgRNA transfer plasmid, 1 μg psPAX2, and 0.1 μg pMD2.G were diluted in 200 μL Opti-MEM I reduced-serum media and combined with 6 ul transIT. Following a 10 minute incubation, this mixture was added dropwise to confluent HEK-293T cells, with lentivirus collected as noted above.

### CRISPRi cell line construction

The dCa9-KRAB and sgRNAs constructs were integrated into uHL-60 cells using a lentivirus spinoculation protocol. Briefly, lentivirus was added to 1 mL cells (1x10^6^ cells /mL) and polybrene reagent (final concentration of 1 μg/mL) in 24-well tissue culture plates. Cells were spun at 1,000 g for two hours at 33°C. Virus was removed and cells were placed in an incubator for two days prior to antibiotic selection for 6 days (dCas9-KRAB: blasticidin 10 μg/mL; sgRNA constructs: puromycin 1 μg/mL).

For CRISPRi sgRNA library preparations, lentiviral titers were estimated by titrating lentivirus over a range of volumes (0 μL, 75 μL, 150 μL, 300 μL, 500 μL, and 800 μL) with 1x10^6^ cells in a total of 1 mL per well of a 24-well tissue culture plate, using the spinoculation protocol noted above. Two days post-transduction, cells were split into two groups, with one placed under puromycin selection. After 5 days, cells were counted for viability. A viral dose that led to a 12.5% transduction efficiency was used for subsequent pooled library work. This low efficiency was targeted to ensure most cells only received one sgRNA integration ^9^. For library work, roughly 230 million cells were transduced, targeting a final number of roughly 30 million successful sgRNA integrations following puromycin selection (or about 500 cells per sgRNA).

Cells were maintained across multiple T175 tissue culture flasks with 35 mls of media.

### Genome-wide CRISPRi assays

#### Overview of cell collection and experimental replicates

For CRISPRi proliferation drop-out screens, genomic DNA (gDNA) was collected from uHL-60 cells at two time points, separated by 6-days of proliferation in T175 tissue culture flasks. Each cell preparation was pelleted by centrifugation and frozen for later genomic DNA isolation.

Results from the proliferation screen represent averages across four independently prepared genome-wide CRISPRi libraries. For CRISPRi differentiation drop-out screens, gDNA was collected in uHL-60 and in dHL-60 cells 5-days following the initiation of differentiation in 15 cm tissue culture dishes. Results from differentiation screens represent averages across four independently prepared genome-wide CRISPRi libraries, each performed twice (8 experiments total).

For cell migration experiments, gDNA was collected from three populations: On the day of the experiment, 5-day differentiated dHL-60 cells were collected, with 3x10^7^ cells set aside for our reference sgRNA population. For chemotaxis and chemokinesis experiments involving track-etch membranes, the other two populations were the fraction of cells that migrated through the membrane (4ii in Fig. 1C) and the fraction of cells that remained on top of the membrane (4i in Fig. 1B). For the amoeboid 3D screen, the other two populations were the fraction of cells that migrated into the fibrin (5ii in Fig. 1C) and the fraction of cells that remained in collagen (5i in Fig. 1C).

All cell migration experiments utilized 5 day differentiated dHL-60 cells with 3x10^7^ cells as reference controls. Chemotaxis and chemokinesis cell migration experiments with track-etch membranes compared the number of cells that migrated through the pores (4ii in Fig. 1C) to those that did not (4i in Fig. 1B), with respect to the reference sample. For the 3D amoeboid migration screen we examined the fraction of cells that migrated into the fibrin (5ii in Fig.1C) compared to those that remained in the collagen (5i in Fig. 1C).

For assays using track-etch membranes, 6-hour time point chemotaxis experiments were performed across four independently prepared genome-wide CRISPRi libraries, with 8 experiments in total. For 6-hour chemokinesis experiments and all 2-hour time points (both, chemotaxis and chemokinesis), experiments were performed using two independently prepared genome-wide CRISPRi libraries (two experiments each). For amoeboid experiments, results represent the average across two independently prepared genome-wide CRISPRi libraries, each performed 3 times (6 experiments total).

#### Removal of cell debris and dead cells prior to cell migration assays

Cellular debris and dead cells were removed from the dHL-60 cell suspensions using density gradient centrifugation. Pooled CRISPRi libraries were differentiated in 15 cm dishes (55 ml cell culture per dish) and eight plates were combined for a single preparation. Cells were first spun down (10 minutes at 300 g), resuspended in 10 mL PolymorphPrep (Cosmo Bio USA #AXS1114683) and added to the bottom of a 50 mL conical tube. Using a transfer pipette, 15 mL of 3:1 PolymorphPrep : RPMI media + 10% hiFBS was gently layered on top by dispensing along the walls of the tube. This was followed by layering another 14 mL of RPMI media + 10% hiFBS. Cells were centrifuged at 700 g for 30 min with the centrifuge acceleration and brake set to half-speed. Live dHL-60 cells were collected between the RPMI media and the 3:1 PolymorphPrep. RPMI media layers were diluted with one volume of RPMI media + 10% hiFBS, and spun down once more for 10 minutes at 300 g. Finally, cells were resuspended in 10 ml RPMI media and counted using a BD Accuri C6 flow cytometer (live cells identified by their forward-scatter and side-scatter).

### Cell migration assays: Chemotaxis and chemokinesis using track-etch membranes

Chemotaxis and chemokinesis transwell migration assays used track-etch membranes with 3 µm pore sizes (6-well plates with 24.5 mm diameter inserts; Corning, #3414). For each experiment, 24 million cells were distributed across four 6-well plates. For each track-etch membrane insert, one million cells were diluted in 1.5 mL media and added to the top of the track-etch membrane. Note that for chemotaxis experiments, cells were purified and resuspended in RPMI media without hiFBS. To the bottom reservoir, 2.6 mL RPMI media with 10% hiFBS was added and the plates were carefully moved to a 37°C incubator. Following incubation for the required time (2 hour and 6 hour time points), the track-etch inserts were separated from the bottom reservoir. To ensure more complete recovery of the migratory cells, the bottom side of the track-etch membrane was gently scraped using a cell-scraper (Celltreat, #229310) to dislodge any cells remaining on the membrane surface. The migratory cells (bottom reservoir) and remaining cells (top reservoir) were separately collected by centrifugation. Following an additional wash with 1 mL PBS, cells were pelleted and frozen at -80°C for later genomic DNA extraction.

Note that in a second set of chemotaxis experiments (6-hour time point; four of the replicates), custom devices were fabricated to house larger, 49 mm track-etch membranes (3 µm pore size, Sigma #TSTP04700). Similar cell densities were targeted as the experiment using multi-well plates above.

#### Cell migration assays: Amoeboid 3D using collagen/ fibrin ECM

Cells were seeded into a multi-layer system of collagen (rat-tail collagen used throughout; ThermoFisher # A1048301) and fibrin as shown in Fig. 1C. Briefly, these were prepared by first creating a ∼50 μm layer of collagen on top of 25 mm glass coverslips, seeding dHL-60 cells in another layer of collagen ∼150 μm thick, and then overlaying this with fibrin ECM. Each genome-wide CRISPRi screen involved our pooled dHL-60 library spread across 32 coverslips with 1x10^6^ cells added to each coverslip.

The glass coverslips were first surface modified to support an adhered layer of collagen ECM using a silane treatment ^124^. A 2% aminosilane solution was first prepared in 95% ethanol/ 5% water and incubated for 5 minutes to allow silanol formation. Coverslips were then immersed in the solution for 10 minutes, rinsed with 100% ethanol, and then cured on a hot plate heated to 110°C for 5-10 minutes. The coverslips were then immersed in 0.25% glutaraldehyde for 15 minutes and then rinsed in water for five minutes. This 5 minute wash was repeated two more times. We then prepared the initial collagen layer by pipetting 22.6 uL of a 1.5 mg/mL collagen mixture (for 3 ml: 1.5 mL 3 mg/mL collagen, 376 μL 0.1M NaOH, 210 μL 10x PBS, and 923 μL PBS) onto a 15 cm plastic tissue culture dish and then placing a coverslip on top, causing the mixture to spread across the entire coverslip. Approximately 16 coverslips were prepared inside a single 15 cm tissue culture dish. The dish was placed in an incubator at 37°C for 18 minutes to gel.

Coverslips containing the initial layer of collagen were carefully removed using tweezers and flipped collagen side up to allow dHL-60 cells to be seeded. Here, a 1 mg/mL collagen mixture was prepared and mixed with dHL-60 cells (recipe for 1.2 ml: 400 μL 3 mg/mL collagen, 100 μL 0.1M NaOH, 55 μL 10x PBS, 525 μL PBS, and 120 μL hiFBS). For each collagen treated coverslip, 3 x 11.3 μL aliquots were pipetted on top of the initial collagen layer, which wetted and spread across the initial collagen layer. Note that the initial coverslip and collagen layer will begin to dry out while inside a biosafety cabinet due to the air flow, so this second collagen mixture must be added relatively quickly to ensure proper spreading of this second layer. The coverslips were again moved to an incubator at 37°C for 15 minutes to gel, placed inside a closed tissue culture dish containing a wetted kimwipe to minimize evaporation. Note that during this time, prior to gelling, cells will settle down to the initial collagen layer.

The final layer, composed of fibrin ECM ^125^, was then prepared on top of the collagen. Here, a 1 mg/mL fibrin ECM was generated by mixing 1 μL thrombin (100 U/ml, Sigma-Aldrich #T1063-250UN) per 1 ml fibrinogen at 1.5 mg/mL (plasminogen-Depleted from human plasma, Sigma-Aldrich #341578) in 1x Hanks’ buffered salt solution (HBSS; ThermoFisher Scientific, Gibco #14-065-056). This was carefully added on top of the collagen by slowly pipetting near the edge of the coverslip. The fibrin ECM will begin to gel immediately, but the coverslips were further incubated at 37°C for 18 minutes to complete the gelling process. Finally, the multi-layered gels were covered with RPMI media containing 10% hiFBS and incubated for 9 hours for dHL-60 cells to migrate.

Recovery of the most migratory dHL-60 cells from the fibrin ECM were obtained by first incubating coverslip/gels with nattokinase, which specifically degrades fibrin ^33^. Here, the RPMI media was first aspirated from the 15 cm dish and 25 ml PBS containing 2.1 mg/mL nattokinase (Japan Bio Science Laboratory USA inc.) and 0.5 M EDTA were added, incubating at 37°C for 40 minutes. The coverslips and remaining collagen layer was then removed, allowing recovery of the released dHL-60 cells. The remaining coverslips + collagen were rinsed with 25 ml PBS + 10% hiFBS prior to scraping the collagen together for later gDNA extraction.

#### Quantification of sgRNA from CRISPRi libraries

Genomic DNA (gDNA) was isolated using QIAamp DNA Blood Maxi (3x10^7^ - 1x10^8^ cells) or Midi (5x10^6^ - 3x10^7^ cells) kits following protocol directions (Qiagen, #51192 and #51183). gDNA precipitation was then used to concentrate the DNA. Briefly, salt concentration was adjusted to a 0.3 M concentration of ammonium acetate, pH 5.2 and 0.7 volumes of isopropanol were added. Samples were centrifuged for 15 minutes at 12,500 g, 4°C. Following a decant of the supernatant, the gDNA was washed with 10 mL 70% ethanol and spun at 12,500 g for 10 minutes, 4°C. The samples were washed in another 750 μL 70% ethanol, spun at 12,500 g for 10 minutes, 4°C, and decanted. The pellets were allowed to air-dry prior to resuspending them in water. The gDNA concentrations and purity were determined by UV spectroscopy.

The sgRNA sequences from each gDNA sample were PCR amplified for sequencing following protocols provided by the Broad Institute’s Genetic Perturbation Platform. Briefly, gDNA samples were split across multiple PCR reactions, with 10 μg gDNA added per 100 μL reaction: 10 μL 10x Titanium Taq PCR buffer, 8 μL dNTP, 5 μL DMSO, 0.5 μL 100 μM P5 Illumina sequencing primer, 10 μL 5 μM P7 barcodedIllumina sequencing primer, and 1.5 μL Titanium Taq polymerase (Takara,# 639242). The following thermocycler conditions were used: 95°C (5 minutes), 28 rounds of 95°C (30 s) - 53°C (30 s) - 72°C (20 s), and a final elongation at 72°C for 10 minutes. PCR products (expected size of ∼360 bp) were gel extracted using the QIAquick gel extraction kit (Qiagen, #28704) following protocol directions. After elution, samples were further cleaned up using isopropanol precipitation. Here, 50 μL PCR DNA samples were combined with 4 μL 5M NaCl, 1 μL GlycoBlue coprecipitant (ThermoFisher Scientific Technologies,# AM9515), and 55 μL isopropanol. Samples were incubated for 30 minutes and then centrifuged at 15,000 g for 30 minutes. The resulting pellet was washed twice with 70% ice-cold ethanol and resuspended in 25 μL of Tris-EDTA buffer. Illumina 150bp paired-end sequencing was performed by Novogene Corp. (Sacramento, CA).

Sequence reads were quality filtered by removal of reads with poor sequencing quality, and reads were associated back to their initial samples based on an 8 bp barcode sequence included in the P7 PCR primer. The 20 bp sgRNA sequences were identified and mapped to gene targets using a reference file for the genome-wide CRISPRi library ^9^.

Log_2_ fold-change values were calculated from the sequencing counts between two sets of samples. The specific comparison sets made for each screen are described in section, ‘Overview of cell collection and experimental replicates’. Note that each cell migration assay resulted in two log_2_ fold-changes measurements, since two populations of cells were collected and each compared to a reference set of sgRNA. Since we expect these two measurements to be inversely correlated, the log_2_ fold-change values from the less-migratory population were multiplied by -1. For example, if gene knockdown resulted in a negative log_2_ fold-change in the bottom reservoir of our chemotaxis track-etch membrane experiment, we would expect a positive log_2_ fold-change for that sgRNA in the upper reservoir. The multiplication by -1 allowed the two sets of log_2_ fold-change values to be compared directly and we averaged across all such measurements.

Reported Log_2_ fold-changes represent averages across median-normalized replicate measurements from the multiple experiments performed. Here, a pseudocount of 32 was added to the sgRNA counts to minimize erroneously large fold-change values in cases of low library representation ^126^. For the differentiation screen and cell migration screens, log_2_ fold-changes were also scaled to have unit variance prior to averaging across individual experiments ^127^. p-values were determined by performing permutation tests ^128^ between the calculated fold-change values for each gene target and our set of ∼500 control sgRNA fold-change values. Adjusted p-values were determined using the Benjamini–Hochberg procedure ^129^.

### Gene set enrichment analysis

Enrichment analysis was performed using the GSEA software version. 4.1.0 ^130^ following recommended parameters ^35^. A subset of significant gene ontology terms are shown.

### Image acquisition

All microscopy-based image acquisition was performed using setups operated by MicroManager 2.0 ^131^.

Cell Migration assays using individual sgRNA CRISPRi lines

#### 2D migration on fibronectin coverslips

Ibidi µ-Slide I slides (#80106, IbiTreat) were incubated with 10 μg/mL fibronectin (from human plasma, #2006, Sigma-Aldrich Inc.) in PBS at room temperature for 1.5 hr. The channel slides were then washed once with 1 mL RPMI media containing 10% hiFBS, once with Leibovitz’s L-15 Medium (ThermoFisher Scientific, Gibco #11415064) , and then the media was removed from the reservoirs of the channel slide. Separately, 2x10^5^ dHL-60 cells were collected, resuspended in 1 mL L-15 + 10% hiFBS media containing 1 μg/mL DNA stain Hoechst 33324 and incubated at 37°C for ∼15 min. Cells were then spun down, resuspended in 200 μL L-15 + 10% hiFBS media, and pipetted into one side of the channel slide inlet. Following a 30 minute incubation at 37°C to allow cells to settle and adhere, the slide was rinsed with three washes of L-15 + 10% hiFBS media to remove any remaining floating cells.

Cells were imaged for nuclei tracking at 37°C on an inverted microscope (Nikon Ti Eclipse), using a 20x objective lens (Nikon 20x 0.75 NA plan apo phase contrast), with sequential phase contrast and epifluorescence illumination through a standard DAPI filter set. For each sample, a 30 min time-lapse movie was acquired with 60 s intervals. Individual experiments were performed over 2-3 days, with five different fields of view during each acquisition. For the higher-magnification still images, cells were imaged with a 100x objective lens (Nikon 100x 1.45NA plan apo) with an additional 1.5x intermediate magnification. Images were captured on an iXon EMCCD camera (Andor).

#### 2D migration using agarose overlap and fMLP photo-uncaging

To assess chemotaxis, we stimulated dHL-60 cells with N-formyl-methionine-alanine-phenylalanine (fMLP) as previously described ^30^. Briefly, to reduce adhesion, glass bottom 96 well plates (Cellvis, #P96-1.5H-N) were treated with 1% BSA (Millipore Sigma, #A7979) in water for 15 min followed by two washes with water then dried overnight at 37°C incubator with lid slightly ajar. Approximately 1,000 dHL-60 cells labeled with Celltracker Orange (ThermoFisher, #C2927) were plated in each well, in a 5 µl drop of modified L-15 + 2% hiFBS media. Cells were allowed to adhere to the glass for 5 min, before a 195 µl layer of 1.5% low-melt agarose solution in L-15 (Goldbio, #A-204-25) was mixed with L-15 + 10% hiFBS media and overlaid on top. The agarose was allowed to solidify at room temperature for 40 minutes and then the plate was transferred to the 37°C microscope incubator 40 minutes prior to imaging. Imaging is done using a Nikon Ti-E inverted microscope controlled by MATLAB via Micromanager, allowing simultaneous automated imaging of multiple wells in groups. An environmental chamber was used to maintain a 37°C temperature, and wells were imaged at 4X magnification every 30 s using epifluorescence illumination with the X-Cite XLED1 LED (Excelitas Technologies, GYX module). The excitation light was filtered using a Chroma ZET561/10 band pass filter (custom ZET561/640x dual laser clean-up filter). A caged UV-sensitive derivative N-nitroveratryl derivative (Nv-fMLP) of fMLP was used at 300 nM final concentration and uncaged on the microscope by exposure to UV light with a filter cube with 350/50 bandpass filter (max intensity around 360-365 nm). The initial gradient was generated with a 1.5 s exposure of UV light and recharged with 100 ms exposure after every frame. Image processing and statistical analyses of chemotaxis were performed using custom MATLAB software (Collins et al., 2015). Note that for each experiment, each cell line was added to 24-32 wells. Each well contained roughly 100 cells, resulting in about 20,000 cells quantified for each cell line across five experiments.

#### 3D migration in collagen ECM

Cells were prepared as previously described ^123^. Briefly, 2x10^5^ dHL-60 cells were collected, resuspended in 1 mL L-15 + 10% hiFBS media containing 1 μg/mL DNA stain Hoechst 33324 and incubated at 37°C for ∼15 min. During incubation with Hoechst stain, a 200 μL collagen aliquot was prepared: 6.5 μL 10x PBS, 12.5 μL 0.1 M NaOH, 111 μL L-15, and 20 μL hiFBS were combined with 50 μl 3 mg/mL collagen. The cell suspension was spun down and resuspended in the collagen mixture for a final concentration of 0.75 mg/mL collagen, and then added to the channel of an Ibidi μ-Slide I (Ibidi, #80106). After 1 min incubation at room temperature, the channel slide was inverted to help prevent cell sedimentation and incubated at 37°C for gel formation. After 20 min, the channel slide media reservoirs were filled with 2 ml total L-15 media containing 10% hiFBS and imaged within 30 minutes to 1.5 hr.

Cells were imaged at 37°C on an inverted microscope (Nikon Ti Eclipse) with a 20x 0.75 NA objective lens using sequential phase contrast and epifluorescence illumination through a standard DAPI filter set. For each sample, a 60 min time-lapse movie was acquired at 60 s intervals. A z-stack was acquired over 200 μm with acquisitions every 3 μm. In general experiments were performed over three different days, with two 60 minute acquisitions taken each day. Images were captured on an iXon EMCCD camera (Andor).

### Cell Tracking and Quantification of Cell Migration Characteristics

Cell tracks were extracted from the DNA channel of time-lapse microscopy images using custom Python code ^123^. Briefly, nuclei were first identified using a morphological mean filter with a 50 pixel radial disk structural element and thresholding using the Python package scikit-image ^132^. For each nuclei identified, the z-coordinate was calculated by taking a weighted-intensity average along the z-axis. With cell coordinates in hand, cell trajectories were determined by calculating all possible cell-to-cell displacements between consecutive time points, and then matching cells through minimization of the total displacement across cells. For example, cells whose displacement changed very little between two time points would most likely correspond to the same cell. The cell density in the ECM was kept low enough that individual cell tracks could be easily identified.

Non-overlapping velocities and cell speeds were calculated using the 30 s (fibronectin-coated coverslips) and 60 s (collagen ECM) frame rate of our image acquisition. To estimate average migratory persistence from each cell trajectory, cell tracks were analyzed using a Bayesian inference algorithm based on a persistence random walk ^59^. Specific parameters were chosen empirically to best capture persistence changes in the tracks (inference grid size = 200, pMin = 10^-5,^ persistence box kernel radius = 2, activity box kernel radius= 2). Persistence were allowed to range from –1.5 to 1.5, and activities were allowed to range from 0 to 0.5 μm/sec.

### Immunolabeling for Flow cytometry

Live-cell immunofluorescence measurements of cell surface expression for CD11b (integrin ɑ_M_), CD18 (integrin ꞵ_2_) and formyl peptide receptor FPR1 were performed on a Sony SH800 Cell Sorter. All staining and washes were done with cells suspended in phosphate-buffered saline (PBS) containing 2% hiFBS and 0.1% sodium azide, chilled on ice. For each sample, one million cells were first resuspended in 100 μL buffer containing 5 μL Fc Receptor Blocking Solution (Biolegend, #422302) and incubated for 15 minutes. Cells were then spun down and resuspended in 100 μL of buffer containing fluorescently conjugated antibodies for one hour (5 μL of each antibody per sample). Following staining, the samples were washed three times by resuspending in 300 μL of fresh buffer. Following collection of flow cytometry data, .fcs files were exported and processed using the Python package FlowCytometryTools (version 0.5.0)^133^. Gating was performed on the forward and side scatter to isolate the population of live cells.

#### Antibodies and dilution information

Fluorophore-conjugated anti-human primary antibodies: BB515 Mouse Anti-Human CD11b (1:20; BD Biosciences, #B564517), CD18 Mouse anti-Human, FITC (1:20; BD Biosciences, #B555923), and fMLP Receptor Mouse anti-Human, Alexa Fluor 647(1:20; BD Biosciences, #565623). Isotype controls: BB515 anti-IgG1, (1:20; BD Biosciences, #B564416), FITC anti-IgG1 (BD Biosciences, #B550616), and Alexa Fluor 647 anti-IgG1 (BD Biosciences, #B557714).

### Immunolabeling for Western blots

Whole-cell protein lysates were collected from dHL-60 cells for Western blot analysis. For each sample, 5x10^6^ cells were collected, washed twice in ice-cold PBS, and resuspended in 100 μL RIPA lysis buffer (Cell Signaling, #9806) containing a protease and phosphatase inhibitor cocktail (Cell Signaling, #5872). The suspension was incubated on ice for 10 minutes and vortexed briefly prior to sonication with a bath type sonicator (Diagenode, #B01020001). Sonication was performed on their high power setting at 4°C with five cycles of 30 seconds on and 30 seconds off. Following sonication the suspension was spun at 15,000 g for 10 minutes at 4°C. Finally each sample was diluted with 4x Laemmli SDS-PAGE sample buffer and heated to 98°C for 5 minutes.

Samples were run on 7.5% polyacrylamide gels and and transferred to nitrocellulose membranes (Bio-rad, #1620233) by semi-dry transfer in buffer 10mM CAPS pH 11, 10% methanol. Transferred protein was assayed using a reversible total protein stain kit (Pierce, #24580) prior to blocking in Tris-buffered saline with 0.1% Tween 20 detergent (TBST) with 0.2% fish skin gelatin (FSG) for 30 minutes at room temperature. Protein loading was also assessed by staining the residual protein on the gel using Coomassie stain (0.006% Coomassie R250 with 10% acetic acid). Primary antibodies were diluted in TBST with 0.2% FSG and incubated overnight at 4°C, which were co-stained with an Alexa Fluor 790 Anti-GAPDH antibody for loading control. Blots were washed with TBST for 15 minutes, with buffer exchanged every five minutes, and then stained with an HRP conjugated secondary antibody diluted in TBST with 0.2% FSG. Following incubation for 60 minutes at room temperature, the blots were washed for 30 minutes in TBST, with buffer exchanged every five minutes. The blots were imaged with a digital gel documentation system (Azure c600), allowing for detection of the secondary HRP antibody detected using a chemiluminescence peroxidase substrate kit (Sigma, #CPS-1) and subsequent detection of the GAPDH loading control detected using its laser based infrared detection system. Three to four blots were performed for each measurement.

#### Antibodies and dilution information

Rabbit anti-human primary antibodies: Total S6K (1:1000; Cell Signaling, #2708), p-Thr389 S6K pAb (1:1000; EMD Millipore, #07-018-I), Total Akt (1:1000; Cell Signaling, #4691), p-Ser473 Akt (1:2000; Cell Signaling, #4060), total mTOR (1:1000; Cell Signaling, #2983), p-Ser2448 mTOR (1:1000; Cell Signaling, #5536), Alexa Fluor 790 to GAPDH (1:1000; Abcam loading control, #ab184578). Secondary antibody: HRP-linked anti-rabbit IgG (1:3000; Cell Signaling, #7074S).

### Immunolabeling for Fluorescence Microscopy during 3D migration

Immunofluorescence imaging was performed in cells migrating in 3D collagen gels. We began by creating a thin layer of collagen on 25 mm glass coverslips. We put a coverslip on top of a 30 μL droplet of 0.75 mg/mL collagen mixture in a glass petri dish, which caused the collagen mixture to spread across the coverslip. The collagen was allowed to gel at 37°C for 90 min and then coverslips were lifted off of the dish by adding PBS and gentle nudging with clean forceps. Coverslips were then flipped collagen-side up and allowed to sit in a culture hood until visibly dry. Next, another 0.75 mg/mL collagen solution was prepared and mixed with HL-60 cells to produce a solution of 5,000-10,000 cells/μL. 30 μL of the cell-collagen suspension was pipetted onto the surface of a collagen-coated coverslip and immediately placed into a 37°C incubator for 20min in a covered petri dish.

Cell-laden gels were then fixed and immunostained, with all steps performed at room temperature. Here, a warmed solution containing 4% PFA, 5% sucrose, and PBS for 20 min. After fixation, coverslips were washed twice for 5 min with PBS. Cells were permeabilized with 0.5% Triton-X 100 in PBS for 10 min, washed twice for 5 min in PBS, and then incubated for 30 min with PBS and 0.05% Tween-20. Samples were then blocked using 20% goat serum in PBS with 0.05% Tween-20 for 30 min. Next, cells were immunolabeled with primary antibodies in PBS, 5% goat serum, and 0.05% Tween-20 for 1hr. After incubation with primary antibody, samples were washed three times for 5 min with PBS and 0.05% Tween-20 and then stained with secondary antibodies, phalloidin, and DAPI, diluted in 1x PBS, 5% goat serum, and 0.05% Tween-20. Following fluorescent labeling, samples were washed three times for 5 min with PBS and 0.05% Tween-20 and once for 5 min in PBS. Samples were then stored at 4°C in PBS or immediately imaged in PBS. Imaging was performed using 3D instantaneous structured illumination microscopy (iSIM) using a VisiTech iSIM mounted on a Nikon Ti Eclipse, with a 100x 1.35 NA silicone oil objective (Nikon). Images were captured using dual CMOS cameras (Hamamatsu, ORCA-fusion Gen III).

#### Antibodies and dilution information

Rabbit anti-human CORO1A (1:100, Cell Signaling, #D6K5B), Mouse anti-human FMNL1 (1:100, Santa Cruz Biotechnology, #sc-390466), Alexa Fluor 488 Phalloidin (1:400, Invitrogen, A12379), Goat anti-Rabbit IgG Alexa Fluor 594 (1:1000, Invitrogen, #A-11012), Rabbit anti-Mouse IgG Alexa Fluor 594 (1:1000, Invitrogen, #ab150116). Note: Control experiments using Alexa Fluor 488, 546, and 647 secondary antibodies without primary antibody each resulted in background signals that appeared as small, bright punctae within the cytoplasm of HL-60 cells and were not used for imaging.

### RNA-Seq

Total RNA was isolated 5x10^6^ cells using the RNAeasy Plus Mini kit (Qiagen, #74134). PolyA enrichment, RNA-seq, sequencing (Illumina), and data processing was performed by Novogene Corp. (Sacramento, CA). Sequencing reads were aligned to Homo Sapiens GRCh38/hg38 genomic research using Hisat2 (v2.0.5)^134^. Differential expression analysis of two conditions/groups, two biological replicates per condition, was performed using the DESeq2 package (v1.20.0) ^135^. The resulting P-values were adjusted using the Benjamini and Hochberg’s approach for controlling the false discovery rate. Genes with an adjusted p-value less than 0.05 found by DESeq2 were assigned as differentially expressed. Enrichment analysis was performed using clusterProfiler ^136^ to identify Gene Ontology (GO) and KEGG pathways with gene sets whose expression was significantly enriched by differential expressed genes.

### Data and code availability

All processed data, code, and figure generation scripts are publicly available as a GitHub repository (https://github.com/nbellive/CRISPRi_screen_HL60_pub).

## Acknowledgements

We thank members of the Theriot lab for useful discussions throughout this work. We also thank Alexander Leydon and Jennifer Nemhauser for access to their Bioruptor instrument, and Takato Imaizumi for access to Bio-Rad equipment used to perform Western blots. N.M.B. was supported as a Fellow of the Jane Coffin Childs Memorial Fund for Medical Research and holds a NIH K99/R00 Pathway to Independence award (5K99GM147355). Research support to J.A.T. was provided by the Howard Hughes Medical Institute. S.R.C. is supported by a National Institutes of Health grant (DP2HD094656) and a Sidney Kimmel Foundation Kimmel Scholar Award.

This article is subject to HHMI’s Open Access to Publications policy. HHMI lab heads have previously granted a nonexclusive CC BY 4.0 license to the public and a sublicensable license to HHMI in their research articles. Pursuant to those licenses, the author-accepted manuscript of this article can be made freely available under a CC BY 4.0 license immediately upon publication.

## Author Contributions

Conceptualization: N.M.B., J.A.T.

Methodology: N.M.B., M.J.F.

Investigation: N.M.B., A.P.vL., E.A.

Writing - original draft: N.M.B. and J.A.T.

Writing - review and editing: M.J.F., A.P.vL., E.A., S.R.C.

Visualization: N.M.B.

Supervision: J.A.T., S.R.C.

Funding acquisition: J.A.T., S.R.C.

## Notes

https://github.com/nbellive/CRISPRi_screen_HL60_pub

### Competing Interest Statement

The authors have declared no competing interest.

