## Supplemental Figures and Legends for "Cell migration CRISPRi screens in human neutrophils reveal regulators of context-dependent migration and differentiation state"

### Supplemental Data Tables 1-5

The .csv files contains the $\log_2$ fold-changes in sgRNA abundance for the screens of proliferation (Supplemental Table 1), differentiation (Supplemental Table 2), chemokinesis (Supplemental Table 3), chemotaxis (Supplemental Table 4), and 3D amoeboid cell migration (Supplemental Table 5). The sign for each gene indicates whether knockdown led to an enrichment of the associated sgRNA (positive) or depletion (negative).

### Supplemental Video 1

The .avi file contains phase video microscopy associated with the snapshots of Fig. S4A and Fig. 4C and Figure S4A. Scale bar (10 µm) and acquisition time are identified in the video and apply to both videos, which show migration of dHL-60 cells with a control sgRNA or sgRNA targeting ITGB2 for gene knockdown.

### Supplemental Figures

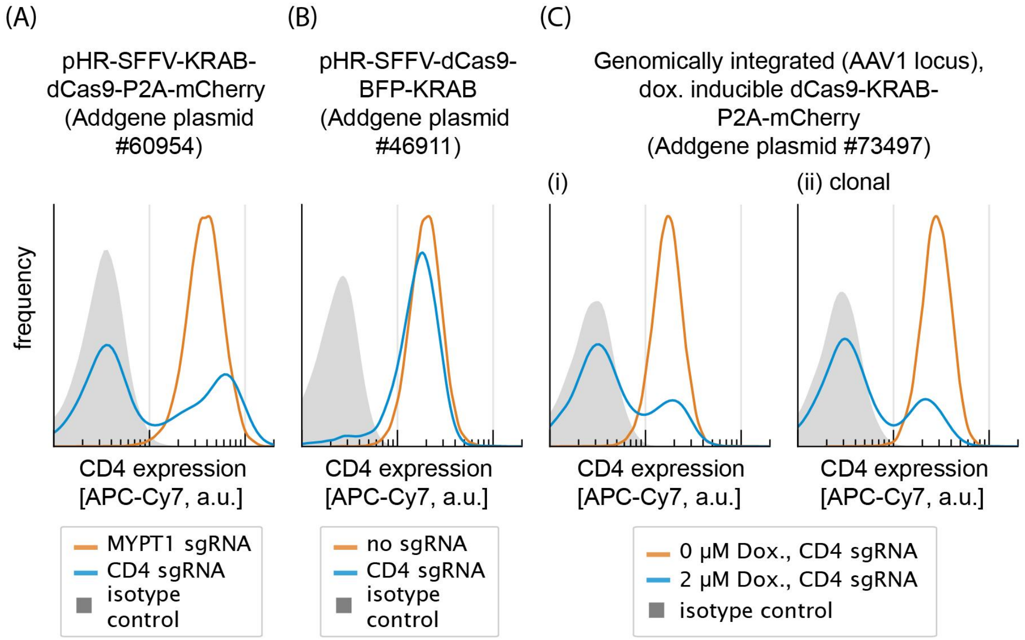


**Figure S1: Comparison of gene knockdown across different dCas9-KRAB constructs. (related to Figure 1)**
Flow cytometry measurements of cell surface CD4 protein expression using three other dCas9-KRAB constructs are shown. Normal expression of CD4 in uHL-60 cells is shown in orange, while cells carrying a sgRNA targeting CD4 are shown in blue. Background autofluorescence and non-specific fluorescence were determined using an isotype antibody control (gray, shaded).

(A) dCas9-KRAB-P2A-mCherry driven by the SFFV promoter (Addgene plasmid #60954). This construct was genomically integrated by lentiviral transduction following the same method described for the construct of the main text.

(B) dCas9-BFP-KRAB (Addgene plasmid #46911). This construct was genomically integrated by lentiviral transduction following the same method described for the construct of the main text.

(C) dCas9-KRAB-P2A-mCherry driven by a doxycycline Inducible promoter (Addgene plasmid #73497). This was genomically integrated into the AAVS1 ‘safe harbor’ locus using co-electroporation with ribonucleoprotein complex of (Cas9 protein and a sgRNA targeting AAVS1) as described in Mandegar et al 2016 (Cell Stem Cell 7;18(4): 541-53).

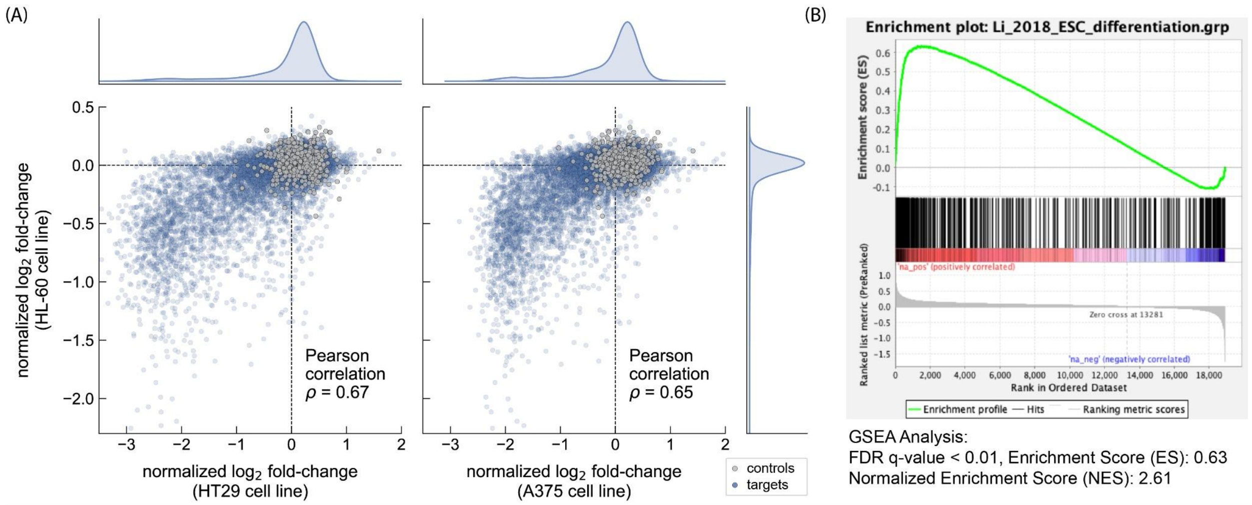


**Figure S2: Additional analysis of cell proliferation and differentiation screens. (related to Figure 2)**

(A) Scatter plots show the normalized $\log_2$ fold-changes measured by comparing the change in sgRNA abundances following six days of growth (HL-60 cell line), plotted against the similar data from Sanson et al. 2018 (HT29, human colorectal adenocarcinoma cell line; A375, human melanoma cell line). Note that the normalized $\log_2$ fold-changes from Sanson et al. 2018 compares sgRNA abundance following 21 days of proliferation, compared to the initial distribution of sgRNAs in the plasmid library preparation.

(B) Gene set enrichment analysis (GSEA) of differentiation data against the genes that were identified in a CRISPR knockout screen for exit from pluripotency in mouse embryonic stem cells (Li et al., 2018). The figure shows an enrichment plot from the GSEA analysis software (Subramanian, Tamayo, et al., 2005) showing a positive enrichment for genes identified in the Li et al. study.

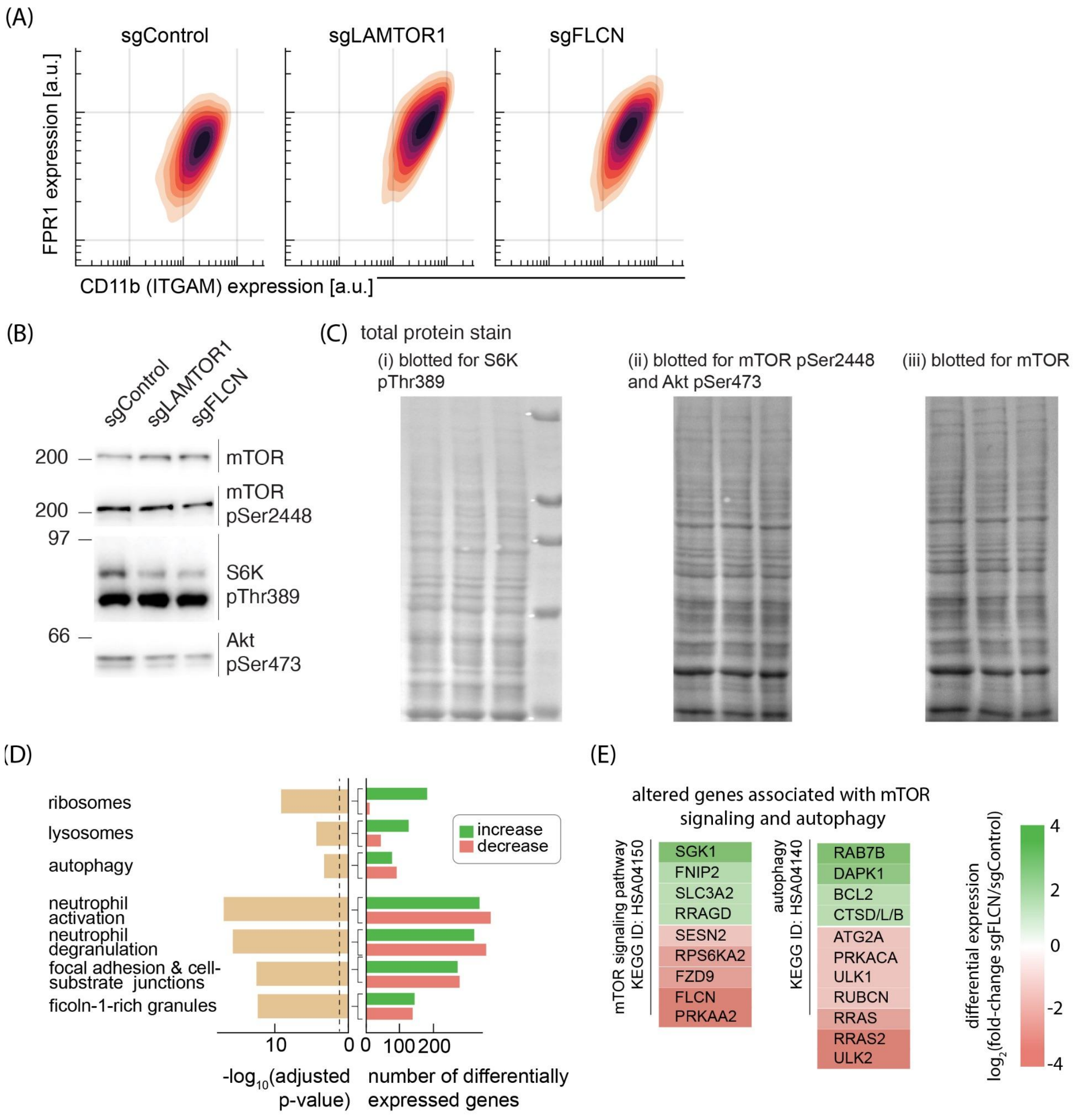


**Figure S3: Comparison of proliferation screen with data from Sanson et al. (related to Figure 3)**

(A) Flow cytometry immunofluorescence measurements of CD11b (ITGAM) and fMLP receptor (FPR1) cell surface expression in dHL-60 cells containing a control sgRNA, LAMTOR sgRNA, and FLCN sgRNA.

(B) Western blots assaying mTORC1/2 activity in differentiated neutrophils (7-day post differentiation) in knockdown lines targeting LAMTOR1, FLCN, and sgRNA control. S6K is a target of mTORC1, while Akt is a target of mTORC2.

(C) Images of total protein stain using a reversible total protein stain, showing uniform loading across samples.

(D) A subset of the pathways identified based on their differential expression between FLCN knockdown cells and control sgRNA (comparison made between day 5 differentiated cells.

(E) Subset of differentially expressed genes ($p_{adj}$ < 0.0001) that were identified and relate to mTOR signaling and autophagy.

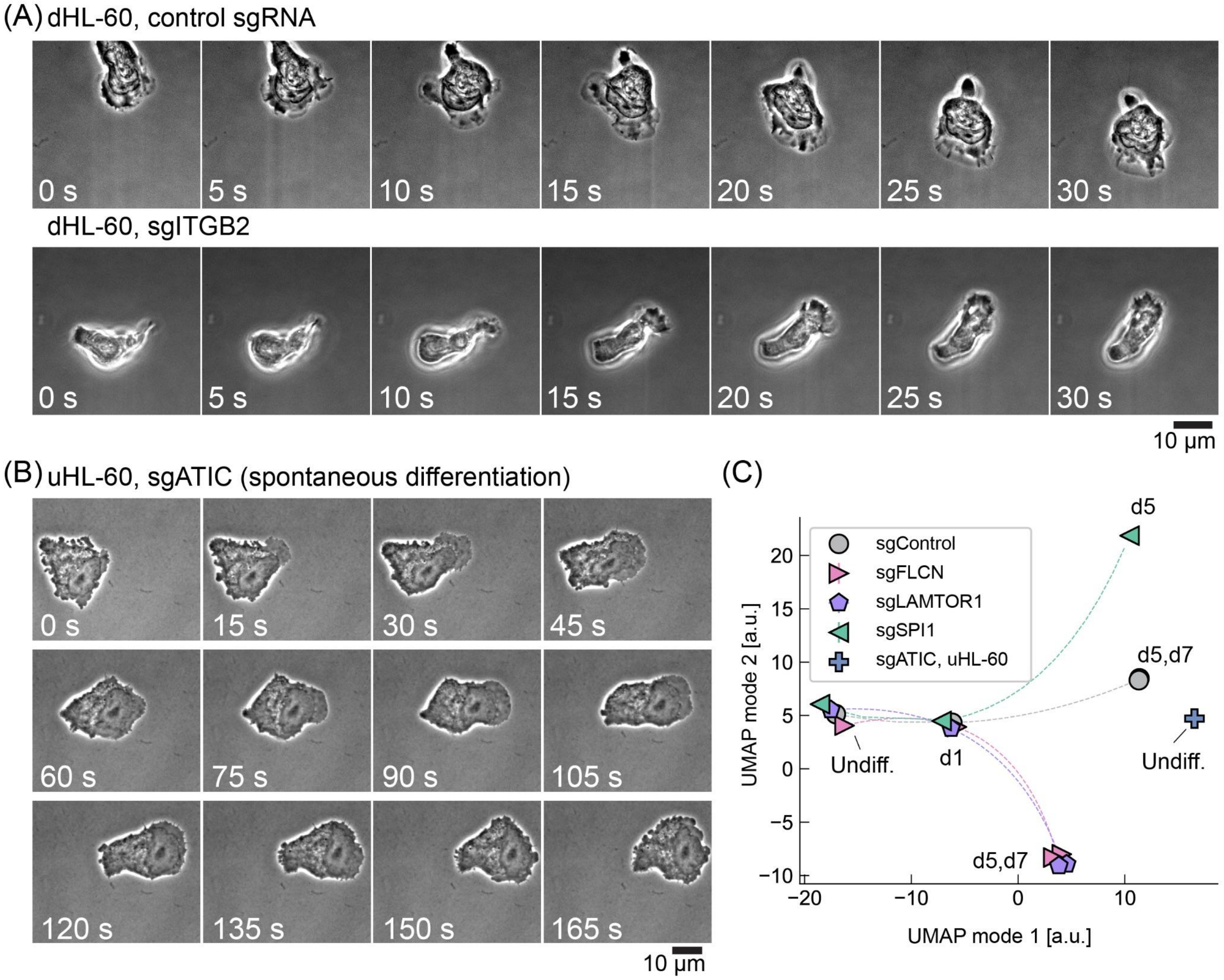


**Figure S4: Additional characterization of cell migration in sgRNA knockdown cell lines. (related to Figure 4)**

(A) Time lapse of phase microscopy images for dHL-60 cells expressing either a control sgRNA (top) or sgRNA targeting ITGB2 for gene knockdown (bottom). Coverslips were coated with fibronectin (see *Methods).* The elongated morphology and phase-bright feature of the ITGB2 knockdown line is because the cell is detached and rounded up, floating above the coverslip.

(B) Time lapse of phase microscopy images of an uHL-60 cell with sgRNA targeting ATIC for gene knockdown. The cell was placed under an agarose overlay in order to confine the cell and prevent movement/detachment due to fluid flow.

(C) The plot shows the addition of uHL-60 cells with sgRNA targeting ATIC to the UMAP dimensionality reduction analysis from the main text (‘+’ symbol), which places them more in line with differentiated dHL-60 cells. Individual data points represent an average across 6 RNA-seq replicate samples.
